# *In vivo* genome-wide CRISPRi-Seq in *Streptococcus pneumoniae* reveals host-adaptive pathways essential for meningitis and identifies new therapeutic targets

**DOI:** 10.1101/2025.10.31.685840

**Authors:** Kin Ki Jim, Andrew Quinn, Vincent de Bakker, Gilles Willemin, Philipp Engel, Nadine Vastenhouw, Jan-Willem Veening

## Abstract

*Streptococcus pneumoniae* is an important human pathogen that causes invasive diseases and remains the leading cause of bacterial meningitis. However, the bacterial determinants required for survival within the central nervous system (CNS) are poorly understood. Here, we performed a genome-wide CRISPR interference screen (CRISPRi-Seq) in an *in vivo* zebrafish model to systematically map pneumococcal fitness determinants during meningitis. Using this approach, we investigated essential and non-essential genes contributing to the pathogenesis of pneumococcal meningitis. The screen identified 244 loci whose repression significantly reduced bacterial fitness *in vivo*, representing pathways involved in virulence, metabolism, cell-envelope biogenesis, translation, and stress adaptation. Comparative analysis with *in vitro* datasets showed that metabolic genes such as *purA, proABC, thrC, glyA,* and *manLMN,* as well as those involved in peptidoglycan and capsule biosynthesis *(pbp3, cps2 operon)* and oxidative stress response (*nox, dpr*), become selectively more essential *in vivo.* Functional validation confirmed that these pathways are critical for virulence, nutrient-dependent growth, and oxidative stress response during meningitis. Untargeted metabolomics of infected zebrafish corroborate several observed bacterial genetic sensitivities in the CNS with altered levels of key nutrients such as glucose and 4-aminobenzoic acid. Integration of the CRISPRi-Seq dataset with antibiotic-target annotations revealed both established and previously unrecognised vulnerabilities, including aminoacyl-tRNA synthetases such as leucyl-tRNA synthetase (LeuRS), whose inhibition by the benzoxaborole epetraborole improved host survival, highlighting its potential as a new therapeutic strategy for multidrug-resistant (MDR) *S. pneumoniae.* Together, these findings demonstrate the power of *in vivo* CRISPRi-Seq to define the pneumococcal essentialome under physiological conditions and reveal that metabolic adaptation, cell-envelope maintenance, and the oxidative stress response are central to bacterial survival in the CNS. This fitness map advances our understanding of pneumococcal adaptation in the CNS and identifies promising targets for developing therapies against *S. pneumoniae*.

## Introduction

*Streptococcus pneumoniae* continues to impose a considerable disease burden, particularly affecting young children and the elderly^1,2^. Pneumococcal meningitis is of notable concern due to its high mortality rate and substantial long-term neurological sequelae in survivors^3,4^. While pneumococcal conjugate vaccines have significantly reduced invasive disease caused by vaccine-covered serotypes, the emergence of non-vaccine serotypes and increasing antibiotic resistance present ongoing challenges^3,5–7^. Notably, macrolide resistance has been rising globally and the World Health Organisation has designated macrolide-resistant *S. pneumoniae* as a priority pathogen, underscoring the need for enhanced surveillance and development of alternative treatment strategies^8^.

Understanding how *S. pneumoniae* adapts to the central nervous system (CNS) is fundamental to enhancing our understanding of pneumococcal meningitis. Successful infection requires the bacterium to overcome host barriers, nutrient limitations, and immune defences while simultaneously adapting its metabolism and virulence strategies to ensure survival and persistence in the host environment^3,6,9^. These adaptations involve significant metabolic shifts, including the ability to scavenge scarce nutrients in the cerebrospinal fluid (CSF) and the regulation of central metabolic pathways to balance energy production and stress resistance^9,10^. Additionally, *S. pneumoniae* must counteract complement activation and phagocytosis by neutrophils and microglia, processes that are tightly linked to oxidative burst and antimicrobial peptide release^6,11–13^. Moreover, pneumococcal factors such as the polysaccharide capsule and surface proteins (e.g. PspA, PspC) play critical roles in evading these host defences^6,12–14^. Elucidating these interactions and adaptations/survival strategies enhances our understanding of meningitis pathogenesis and may reveal novel therapeutic targets, highlighting the need for systematic and genome-wide approaches to identify bacterial determinants of survival and adaptation in the CNS.

Genome-wide functional screening tools, such as transposon sequencing (Tn-seq), have been instrumental in identifying non-essential genes contributing to bacterial fitness under various conditions^15,16^. However, Tn-Seq’s reliance on random insertional mutagenesis limits its ability to probe genes essential for bacterial viability^17^. To overcome this limitation, we developed an inducible CRISPR interference (CRISPRi) system specifically tailored for *Streptococcus pneumoniae*^18,19^. This system employs a catalytically inactive Cas9 (dCas9) to reversibly repress gene expression, enabling systematic investigation of both essential and non-essential genes^20,21^. *In vitro* applications of CRISPRi-Seq have provided valuable insights into pneumococcal biology. For instance, high-throughput phenotyping has identified new essential genes involved in cell wall synthesis, antibiotic resistance, DNA replication, and central metabolism, refining our understanding of bacterial growth requirements^19,22,23^. Additionally, CRISPRi-TnSeq and dual CRISPRi-Seq approaches have mapped genetic interactions, revealing functional redundancies and potential targets for antimicrobial development^22,24^. *In vivo*, CRISPRi-Seq has been utilised to assess gene fitness during influenza A superinfection in a mouse pneumonia model^25^. Studies have identified critical bottlenecks in pneumococcal dissemination and highlighted genes essential for survival in host environments, such as those involved in capsule production and purine biosynthesis^25^. Notably, certain genes deemed essential *in vitro*, like *metK*, were found to be dispensable *in vivo*, underscoring the importance of context-specific gene function analysis^25^.

Here, to our knowledge, we employ CRISPRi-Seq for the first time to investigate the bacterial essentialome in the context of pneumococcal meningitis in an *in vivo* whole animal model. Applying this approach during infection allowed us to identify conditionally essential genes that are not apparent *in vitro*, yielding a more physiologically relevant view of pneumococcal survival and adaptation in the CNS. These insights broaden our understanding of the bacterial processes underlying pneumococcal meningitis and offer a basis for future development of targeted therapies.

## Results

### Genome-wide *in vivo* fitness profiling of Streptococcus pneumoniae in a zebrafish meningitis model using CRISPRi-Seq

To systematically identify pneumococcal genes critical for survival and pathogenesis during meningitis, we used a previously developed doxycycline-inducible CRISPRi system in *S. pneumoniae*, which allows precise temporal control of gene knockdown *in vitro* and *in vivo*^18,19,25^. In this system, both the single guide RNA (sgRNA) and catalytically inactive Cas9 (dCas9) are integrated into the chromosome of the serotype 2, antibiotic-susceptible *S. pneumoniae* D39V strain. Expression of dCas9 is controlled by a doxycycline-inducible promoter (Ptet), while the sgRNA is transcribed from a constitutive promoter (P3) (Figure 1A). Upon doxycycline addition, dCas9 is produced and directed by the sgRNA to its target gene or operon, where it forms a transcriptional roadblock and silences gene expression (Figure 1B). In the absence of induction, target genes remain actively transcribed. Here, we applied this inducible CRISPRi system in our established zebrafish meningitis model to systematically interrogate pneumococcal gene function during infection *in vivo* (Figure 1C).

**Figure 1.**
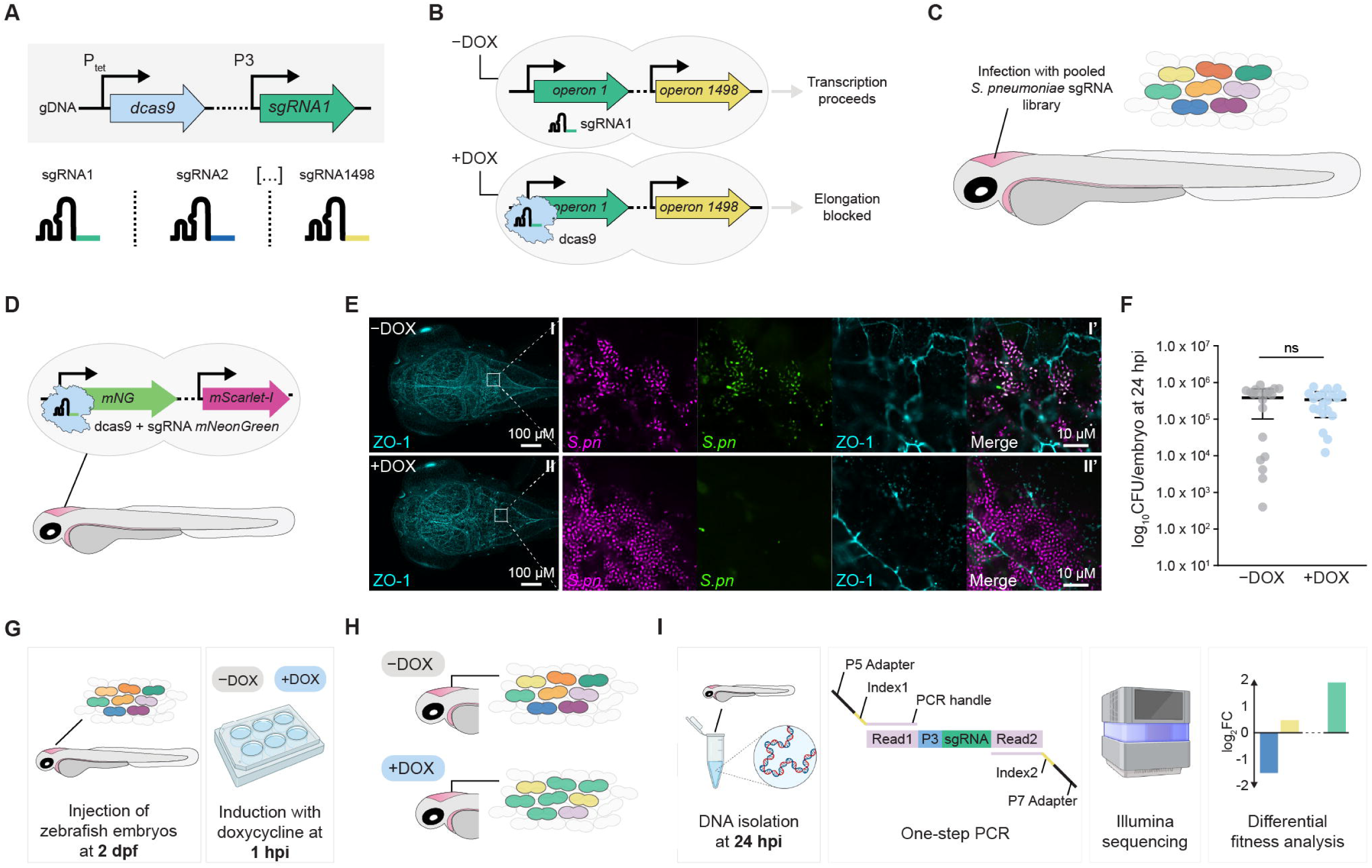
A doxycycline-inducible CRISPRi-Seq system in an *in vivo* zebrafish embryo meningitis model. (A) The two essential CRISPRi components, dCas9 and sgRNA, were integrated into the chromosome of *S. pneumoniae* D39V and placed under control of a doxycycline-inducible promoter (Ptet) and a constitutive promoter (P3), respectively. The pooled CRISPRi library targets all 1,498 operons of the D39V strain (VL4049). (B) Upon addition of doxycycline, dCas9 is expressed and directed by the constitutively produced sgRNA to its specific target sequence, where the dCas9-sgRNA complex binds and blocks transcription of the target gene. In the absence of the inducer, dCas9 is not expressed, and the target gene remains actively transcribed. (C) *In vivo* zebrafish embryos meningitis model used for infection with the *S. pneumoniae* CRISPRi library (VL4049). (D) Reporter strain to assess *in vivo* activity of the doxycycline-inducible CRISPRi system in the zebrafish embryo meningitis model. Strain VL2359 constitutively expresses the fluorescent reporters mNeonGreen and mScarlet-I, with mNeonGreen serving as the CRISPRi target. (E) Zebrafish embryos were infected with ∼15,000 colony-forming units (CFU) of VL2359 at 2 days post-fertilisation (dpf), and either DMSO control (I) or 500 ng/mL doxycycline (II) was added 1 h post-infection. At 24 h post-infection, fixed embryos were imaged by confocal microscopy in the red (mScarlet-I), green (mNeonGreen), and far-red (zona occludens-1) channels to assess doxycycline-dependent repression of mNeonGreen expression. Panels I′ and II′ show magnified views of the indicated regions. (F) Bacterial load in zebrafish embryos infected with ∼15,000 CFU of VL2359 and treated with DMSO or 500 ng/mL doxycycline at 24 h post-infection (hpi). Data represent the mean ± SEM from two independent experiments with 10 larvae per group (n = 20/group). Each dot corresponds to an individual embryo; ns, non-significant (*p*-value = 0.5967); determined by Mann-Whitney U test. (G) Workflow of the *in vivo* CRISPRi-seq screening. Zebrafish embryos were injected with VL4049 at 2 days post-fertilisation (dpf) and induced with doxycycline at 1 hours post-injection (hpi). (H) The CRISPRi library was allowed to grow in zebrafish embryos for 24 hours in the presence (+DOX) or absence (-DOX) of the inducer to compare *in vivo* fitness effects. (I) After 24 hours, embryos were pooled, and genomic DNA (gDNA) was extracted and used as a template for PCR amplification of sgRNA sequences. The forward oligo binds to the Illumina amplicon element read 1 and contains the Illumina P5 adapter sequence, while the reverse oligo binds to read 2 and carries the P7 adapter sequence. Index 1 and Index 2 were incorporated into the forward and reverse oligos, respectively, to enable barcoding of different samples. Amplicons were sequenced in a pooled fashion on an Illumina NovaSeq 6000 using a custom recipe, and fitness costs were determined by differential fitness analysis of sgRNA read counts between induced (+DOX) and uninduced (–DOX) samples.

To test the functionality of the doxycycline-inducible CRISPRi system *in vivo*, we used a fluorescent *S. pneumoniae* reporter strain that constitutively expresses mScarlet and suppresses mNeonGreen upon CRISPRi activation (Figure 1D)^25^. Zebrafish embryos were injected at 2 days post-fertilisation (dpf)^26^, and doxycycline was added to the egg water at 1 hpi. At 24 hpi, we observed that a large proportion of pneumococcal cells in the cerebrospinal fluid (CSF) showed loss of mNeonGreen fluorescence, indicating transcriptional repression *in vivo* at doxycycline concentrations of at least 500 ng/mL (Figure 1E). Importantly, doxycycline at this concentration did not affect bacterial growth in the hindbrain ventricle or *in vitro* (Figure 1F).

For the genome-wide CRISPRi-seq screening of *S. pneumoniae* during meningitis, we injected a pooled *S. pneumoniae* sgRNA library (VL4049) into the hindbrain ventricle of embryos at 2 dpf^25,26^ (Figure 1G). This library comprises 1,498 sgRNAs designed to target 2,111 out of 2,146 annotated genetic elements in the wild-type D39V strain^25^. Infected zebrafish embryos were then treated with doxycycline or vehicle control (DMSO) by supplementing the egg water at 1 hour post-injection (hpi). At defined time-points (e.g. 24 hpi), pooled embryos were collected and bacterial genomic DNA extracted, followed by one-step PCR to amplify sgRNA sequences and subsequent Illumina sequencing (1H-I). The sgRNAs were then quantified from the raw reads, and their abundances between induced and non-induced conditions were compared to determine differences in gene fitness *in vivo* during infection (Figure 1I).

### Identification of genes important for pneumococcal fitness during meningitis

In total, four biological replicates consisting of 50 embryos each were infected with the pooled *S. pneumoniae* sgRNA library for each condition, either doxycycline-treated to induce gene repression or untreated (n = 200 per condition) (Figure 2A). Principal component analysis (PCA) of normalised sgRNA read counts revealed clear and consistent clustering of doxycycline-induced and non-induced conditions, indicating effective transcriptional repression and reproducibility of the *in vivo* screen (Figure 2B). Upon induction, multiple sgRNAs consistently dropped out, indicating that repression of the corresponding genes impaired bacterial fitness during infection (Figure 2C). Moreover, sgRNA abundance profiles were highly consistent across biological replicates, as shown by hierarchical clustering and heatmap analysis, underscoring the reproducibility of the CRISPRi-Seq screen in the zebrafish embryo meningitis model (Figure 2D). Importantly, no signs of a population bottleneck were observed, suggesting that sgRNA diversity was maintained in vivo during infection.

**Figure 2.**
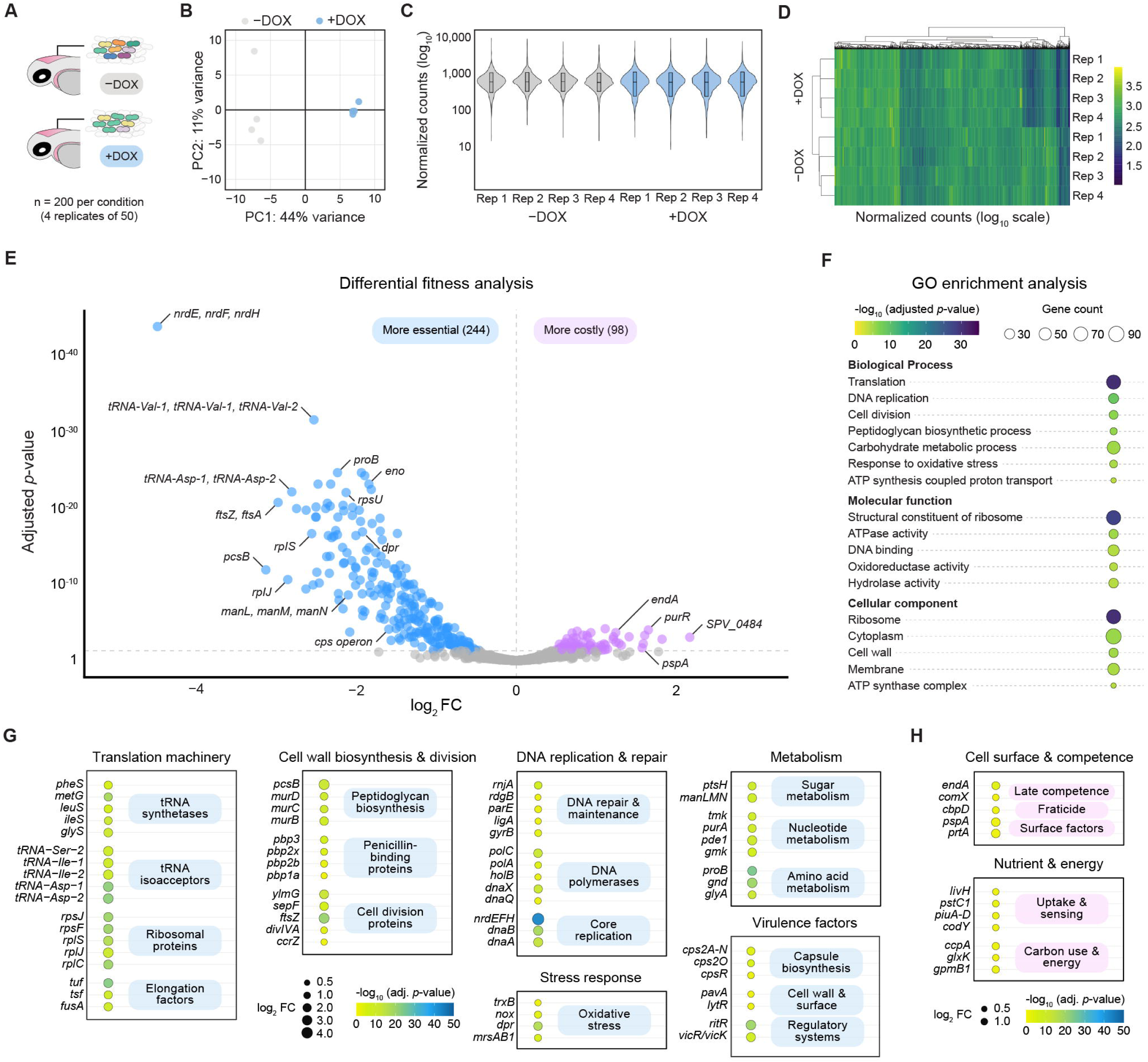
*In vivo* CRISPRi-seq reveals gene fitness determinants during pneumococcal meningitis. (A) Experimental design of the *in vivo* CRISPRi-seq screen. A total of 400 zebrafish embryos were infected with ∼15,000 colony-forming units (CFU) of the *S. pneumoniae* CRISPRi library (VL4049), divided equally between +DOX (500 ng/mL doxycycline) and –DOX (DMSO control) conditions (n = 200 per condition, 50 embryos per replicate, 4 biological replicates). (B) Principal component analysis (PCA) of CRISPRi-seq readouts from *in vivo* zebrafish meningitis samples with (+DOX) and without (–DOX) induction. (C) Violin plots showing the normalised abundance of each sgRNA per sample under +DOX (induced) and –DOX (uninduced) conditions. sgRNA abundance was calculated as 1,498 × (counts of a given sgRNA / total counts of all sgRNAs). Black line = median; box = 10th-90th percentiles. (D) Heatmap of pairwise sample correlations based on sgRNA abundance, visualized with hierarchical clustering, demonstrating high consistency across biological replicates and treatment groups. (E) Volcano plot of sgRNA target fitness scores comparing bacterial growth during meningitis with (+DOX) and without (–DOX) CRISPRi induction. Significance thresholds: FDR-adjusted p < 0.05 and |log₂FC| > 1 (dotted grey lines), calculated with DEseq2. Genes whose repression led to sgRNA depletion were classified as more essential, while enrichment indicated more costly genes under *in vivo* conditions. (F) Gene Ontology (GO) enrichment analysis of genes classified as more essential during pneumococcal meningitis, highlighting overrepresented biological processes, molecular functions, and cellular components. (F) Functional categorisation of a subset of *in vivo* essential genes. Genes were grouped into major functional categories based on their predicted roles. (G) Functional categorisation of a subset of *in vivo* costly genes. Genes were grouped into major functional categories based on their predicted roles.

Differential fitness analysis of the CRISPRi-Seq screen revealed that 244 sgRNA targets were found to be more essential *in vivo* during pneumococcal meningitis at 24 hpi, while 98 sgRNA targets were more costly (adjusted *p*-value < 0.05) (Figure 2E, Table S1A). Gene Ontology (GO) enrichment analysis of the more essential genes revealed an overrepresentation of Biological Process (BP) categories essential for bacterial growth, cell division, cell metabolism, cell maintenance, and stress response (Supplementary Table S2A). The most significantly enriched term was “Translation” (adjusted *p*-value = 2.1E-31), highlighting extensive representation of genes involved in ribosomal structure and function. Processes such as “DNA replication” (adjusted *p*-value = 1.7E-9) and “Cell division” (adjusted *p*-value = 6.0E-07) were also enriched, highlighting genes essential for genome duplication and cytokinesis. Significant enrichment was further observed for “Carbohydrate metabolic process” (adjusted *p*-value = 5.5E-06) and “Peptidoglycan biosynthetic process” (adjusted *p*-value = 1.5E-06), indicating that many genes in the set contribute to energy production and the synthesis of key cell wall components. Additional overrepresented terms included “Response to oxidative stress” (adjusted *p*-value = 2.0E-05) and ATP synthesis coupled proton transport (adjusted *p*-value = 1.2E-04), underscoring the presence of genes involved in stress adaptation and core energy generation, respectively.

In-depth analysis revealed that indeed a large subset of putative essential genes was involved in protein synthesis and RNA metabolism (Figure 2G, Table S1B). This included ribosomal proteins, assembly factors, elongation and initiation factors, numerous tRNA synthetases, and tRNA isoacceptors, highlighting the importance of efficient translation in the CSF. Genes involved in central metabolism, including biosynthesis, nutrient uptake, and energy generation, were more essential *in vivo*, suggesting a dependence on endogenous biosynthesis and energy production under host conditions (Figure 2H). Genes required for the integrity and remodelling of the bacterial cell envelope were also more essential, including peptidoglycan biosynthesis enzymes, penicillin-binding proteins, capsule and teichoic acid synthesis genes, and autolysins. These factors are important for maintaining structural integrity and evading host immunity. To survive oxidative stress in the host, pneumococci relied on redox-balancing enzymes and stress-responsive regulators. Genes involved in DNA repair, ribonucleotide reduction, and cell division were also essential, underlining the need for continued proliferation and genome maintenance *in vivo*. Finally, stress adaptation systems, such as two-component regulators, manganese transporters, and toxin-antitoxin modules, were essential for environmental sensing and maintaining homeostasis *in vivo*. Notably, over 60 SPV-annotated genes with no prior functional data were also identified as essential *in vivo*, suggesting the presence of previously unrecognised mechanisms supporting pneumococcal adaptation and virulence.

In total, 98 sgRNA targets were significantly more costly *in vivo* during pneumococcal meningitis at 24 hpi (adjusted *p*-value < 0.05), indicating that repression of these genes enhanced bacterial survival during infection (Figure 2H, Table S1C). Although Gene Ontology (GO) enrichment analysis did not show statistically significant terms, the costly gene set included hits related to stress response, nutrient transport, and transcriptional regulation (Table S2B). Many encoded nutrient uptake systems that likely become energetically unfavourable or poorly matched to the CSF environment, such as the complete iron ABC transporter (*piuA-D*), amino acid and peptide transporters, and carbohydrate metabolism genes. Their repression may reduce metabolic burden or prevent the accumulation of toxic intermediates under nutrient-limited conditions. Transcriptional regulators, including *codY* and *ccpA*, which coordinate nitrogen and carbon metabolism, were also costly, suggesting that unrestrained biosynthetic activation reduces fitness *in vivo*. Similarly, competence regulators (*comX1*, *comX2*, *ccnE*) may induce non-essential pathways late in infection. Among surface-associated factors, including *pspA*, *cbpD*, and *prtA*, were costly, possibly reflecting increased immune recognition or high biosynthetic cost. Together, these findings suggest that pneumococcal adaptation to the CNS requires not only activation of key survival pathways but also downregulation of energetically demanding or maladaptive processes.

### Interaction analysis identifies pneumococcal genes required for *in vivo* virulence

We next compared *in vivo* fitness data to a previously published *in vitro* CRISPRi-Seq dataset obtained from bacteria grown in C+Y laboratory medium (Figure 3A) to identify genes that became selectively more essential during pneumococcal meningitis ^25^. PCA of normalised sgRNA read counts revealed clear and consistent clustering of doxycycline-induced (+DOX) and non-induced (–DOX) samples under both *in vitro* and *in vivo* conditions, demonstrating effective transcriptional repression and high reproducibility of the CRISPRi-Seq screens (Figure 3B). This interaction analysis identified 49 sgRNAs with significantly different fitness between conditions (adjusted *p*-value < 0.05), including 22 sgRNAs targeting genes that were more essential *in viv*o, likely reflecting bacterial functions required for survival and persistence within the host (Figure 3C, Table S3). The remaining 27 sgRNAs targeted genes that were less essential *in vivo* than *in vitro*, suggesting that certain bacterial functions may be dispensable or even detrimental during infection of the CNS.

**Figure 3.**
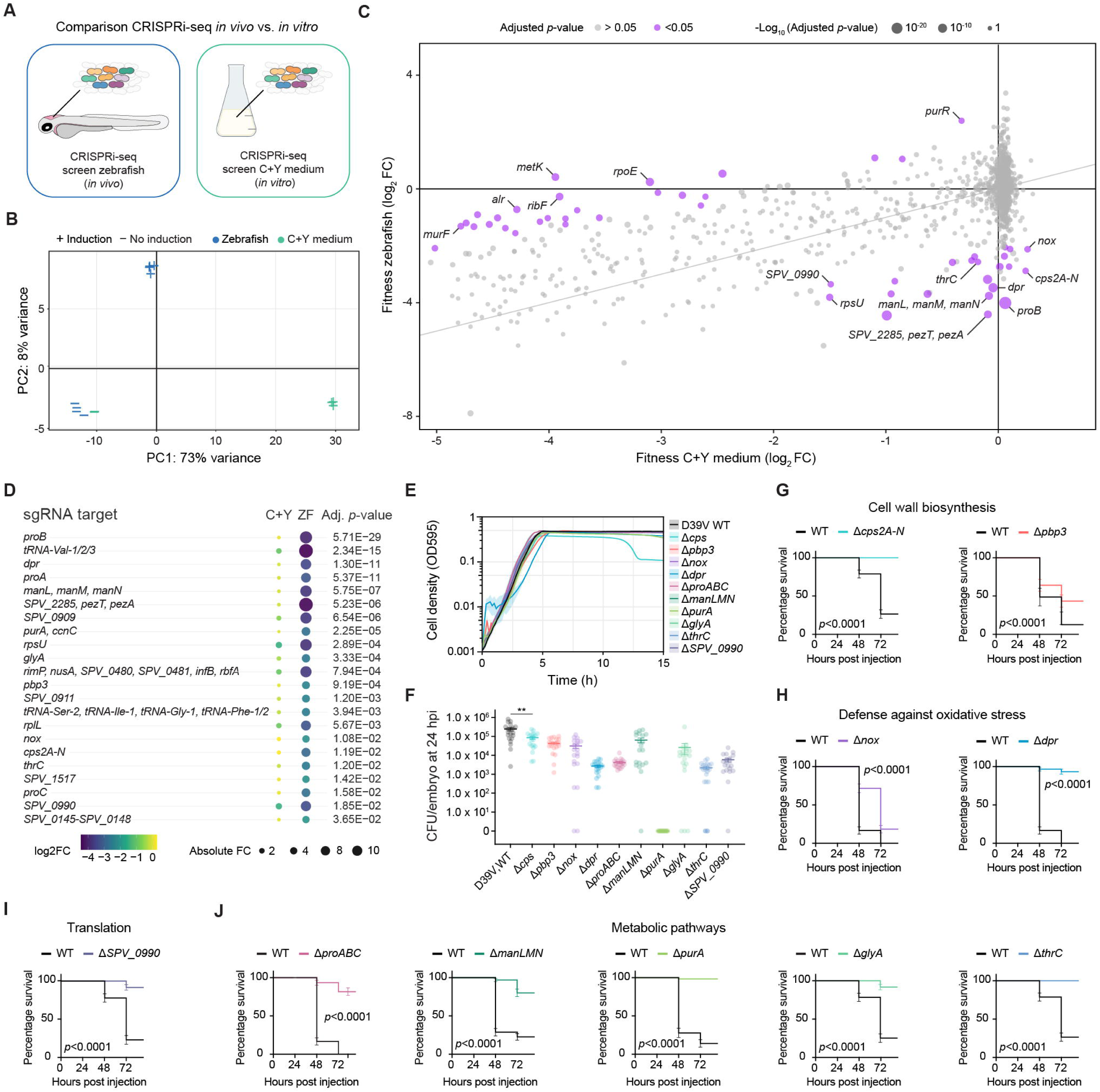
Comparative CRISPRi-seq analysis identifies pneumococcal fitness determinants *in vivo* and validates their roles in meningitis pathogenesis. (A) Schematic overview of the experimental design comparing CRISPRi-seq screens performed *in vivo* in the zebrafish meningitis model and *in vitro* in C+Y medium. (B) Principal component analysis (PCA) of CRISPRi-seq readouts from zebrafish larvae with (+DOX) and without (–DOX) induction, compared to C+Y medium. (C) Differential gene fitness between zebrafish pneumococcal meningitis at 24 hpi and C+Y medium. Values represent log₂ fold changes for the interaction between condition (*in vivo* vs. *in vitro*) and CRISPRi induction (+DOX vs. –DOX), calculated by DESeq2 with *p*-values adjusted by FDR correction. (D) Subset of genes classified as more essential *in vivo* compared to C+Y medium. (E) Growth curves of ten individual knockout mutants that were attenuated in survival experiments and bacterial load experiments. (F) Bacterial load of zebrafish larvae infected with the indicated knockout D39V mutants and D39V wild type strain at 24 hpi. Each dot represents a single larva; data from two independent experiments (n = 20 per group). Bars represent mean ± SEM. Statistical significance was determined using the Mann-Whitney U test (*p*-value = 0.0078; **). (G-J) Survival analysis of zebrafish larvae infected with indicated knockout D39V mutants. Two days post-fertilisation (2 dpf) embryos were injected with 300-400 CFU of pneumococci into the hindbrain ventricle. Fifteen mutants were tested in total; data are shown for the ten that exhibited attenuated virulence *in vivo*. Data represent the mean ± SEM of three biological replicates with 20 larvae per group (n = 60 per condition). Survival data were analysed by the log-rank (Mantel-Cox) test; *p*-value < 0.05 was considered statistically significant. (G) Genes involved in cell-wall biosynthesis (*cps2* operon, *pbp3*). (H) Genes involved in oxidative stress defence (*nox*, *dpr*). (I) Gene involved in translation (*SPV_0990*). (J) Genes involved in metabolic pathways (*proABC*, *manLMN*, *purA*, *glyA*, *thrC*).

To validate the CRISPRi-Seq findings, we selected 17 of the 22 sgRNAs that showed increased essentiality *in vivo* compared to *in vitro* for functional testing in the zebrafish embryos meningitis model (Figure 3D). Three of these targeted different regions of the *proABC* operon, resulting in 15 unique genes or operons chosen for knockout construction. Although sgRNA0005 targets both *purA* and *ccnC*, only purA was deleted, as sgRNA0007 (targeting *ccnC*) showed no induction-dependent dropout or fitness difference between *in vivo* and *in vitro* conditions. For the *pezA-T* toxin-antitoxin system (sgRNA0374), we tested the double mutant because deletion of *pezA* alone was not possible, likely due to toxicity from unopposed *pezT* expression as previously described^25^. *SPV_2285* was excluded, as it represents a pseudogene^27^. Knockout construction was attempted but unsuccessful for genes targeted by sgRNA0184 (*rimP, nusA, SPV_0480, SPV_0481, infB, rbfA*), sgRNA0467 (*rpsU*), and sgRNA0815 (*rplL*), likely reflecting their essential roles in ribosome assembly and protein synthesis. sgRNAs targeting tRNA genes (sgRNA1308, sgRNA1341) were not pursued because these loci are highly conserved, often occur in multiple isoacceptor copies, and are difficult to interpret due to redundancy and potential polar effects^28^.

All successfully constructed deletion strains grew normally in C+Y medium, except for the Δ*dpr* mutant, suggesting that the attenuation observed *in vivo* is most likely not caused by intrinsic growth defects under laboratory conditions (Figure 3E). Among the *in vivo* essential sgRNA targets were genes involved in cell-envelope biogenesis and immune evasion. This included *pbp3* (sgRNA0290), which encodes a penicillin-binding protein required for peptidoglycan biosynthesis, and the *cps2A-cps2N* (sgRNA0127) capsule biosynthesis operon, a major determinant of pneumococcal virulence^12,14^. Deletion of these genes resulted in reduced virulence in survival assays and decreased bacterial burden, underscoring their importance in maintaining structural integrity and avoiding immune detection (Figure 3F and 3G).

Defence against oxidative stress also emerged as an important pathway for pneumococcal survival *in vivo*, with multiple hits targeting genes involved in neutralising reactive oxygen species. Nox (NADH oxidase) catalyses the reduction of O₂ to H₂O₂, while Dpr (a Dps-like peroxide resistance protein) protects against oxidative damage by sequestering iron and limiting Fenton chemistry and are involved in virulence^29,30^. Deletion of *dpr* (sgRNA0525) and *nox* (sgRNA0485) resulted in reduced bacterial loads and attenuated virulence (Figure 3F and 3H). Of note, the attenuated phenotype of Δ*dpr* may partly result from its slower growth rate *in vitro*.

Deletion of *SPV_0990* (sgRNA0809), encoding a putative ribosome-associated protein, also reduced virulence, suggesting that protein synthesis and translational control contribute to metabolic adaptation in the CNS^31^ (Figure 3I). Among the most enriched functional categories were genes involved in amino-acid and nucleotide biosynthesis (Figure 3D). Genes required for proline synthesis (*proA*, *proB*, *proC*; sgRNA1049, sgRNA0314, sgRNA1050), threonine synthesis (*thrC*; sgRNA0697), glycine-serine interconversion (*glyA*; sgRNA1073), and purine biosynthesis (*purA*; sgRNA0005) were significantly more essential *in vivo*. Deletion of these genes led to attenuation in virulence and lower bacterial loads (Figure 3J). Consistent with previous studies, *purA*, encoding adenylosuccinate synthetase, has been linked to pneumococcal virulence in experimental meningitis and pneumonia murine models^25,32^. In addition, manN, manM, and manL (sgRNA0104), encoding components of a mannose-class phosphotransferase system, were significantly more essential *in vivo* than *in vitro*^33,34^. Deletion of this operon impaired bacterial survival in the zebrafish meningitis model, resulting in reduced host mortality and bacterial burden (Figure 3J).

Notably, not all genes classified as more essential *in vivo* were attenuated when deleted and tested experimentally (Figure S1). Mutants lacking *SPV_0909*, *SPV_1517, SPV_0145-0148, ccnC* or the toxin-antitoxin pair *pezT-pezA* displayed survival curves comparable to the wild-type D39V strain (Figure S1A-F). These discrepancies may reflect functional redundancy, limited stress exposure under the tested conditions, or compensatory pathways that mitigate gene loss. Deletion of *SPV_0911* (*staR*) caused a modest, non-significant reduction in survival, with bacterial loads comparable to wild type, suggesting a role in persistence rather than acute replication.

### Metabolic adaptation of *Streptococcus pneumoniae* to the host nutrient environment during meningitis

As shown previously, many genes involved in central metabolism were more essential and their deletion mutants attenuated during pneumococcal meningitis, suggesting that *S. pneumoniae* depends on specific biosynthetic pathways to persist in the nutrient-limited environment of the cerebrospinal fluid (CSF). To test whether these essentialities reflect nutrient limitation, we performed rescue experiments in a CSF-like medium by supplementing the corresponding metabolites^35^. For *purA*, which functions in purine biosynthesis, adenine supplementation fully restored growth but not guanine or xanthine, confirming that purine availability is limiting in CSF-like conditions^25^ (Figure 4A). For the amino-acid biosynthesis genes *thrC* and *proAB*C, we observed partial auxotrophy: both mutants grew without supplementation but showed improved growth when threonine or proline was added, indicating that these pathways confer an advantage when amino acid concentrations are low but not completely absent (Figure 4B and 4C). The Δ*glyA* mutant displayed a different pattern: it grew normally without glycine but poorly without serine, and growth was restored by serine addition, consistent with the role of GlyA in converting glycine into serine and rendering the mutant serine-dependent under nutrient restriction^36^ (Figure 4D). Finally, for the manLMN transporter mutant, both glucose and mannose partially restored growth, but glucosamine provided the strongest rescue, indicating that this phosphotransferase system is flexible but optimised for glucosamine uptake^33^ (Figure 4E). These results demonstrate that the CSF represents a nutrient-poor environment where the limited availability of amino acids, sugars, and purines forces pneumococci to rely on their own biosynthetic and uptake pathways.

**Figure 4.**
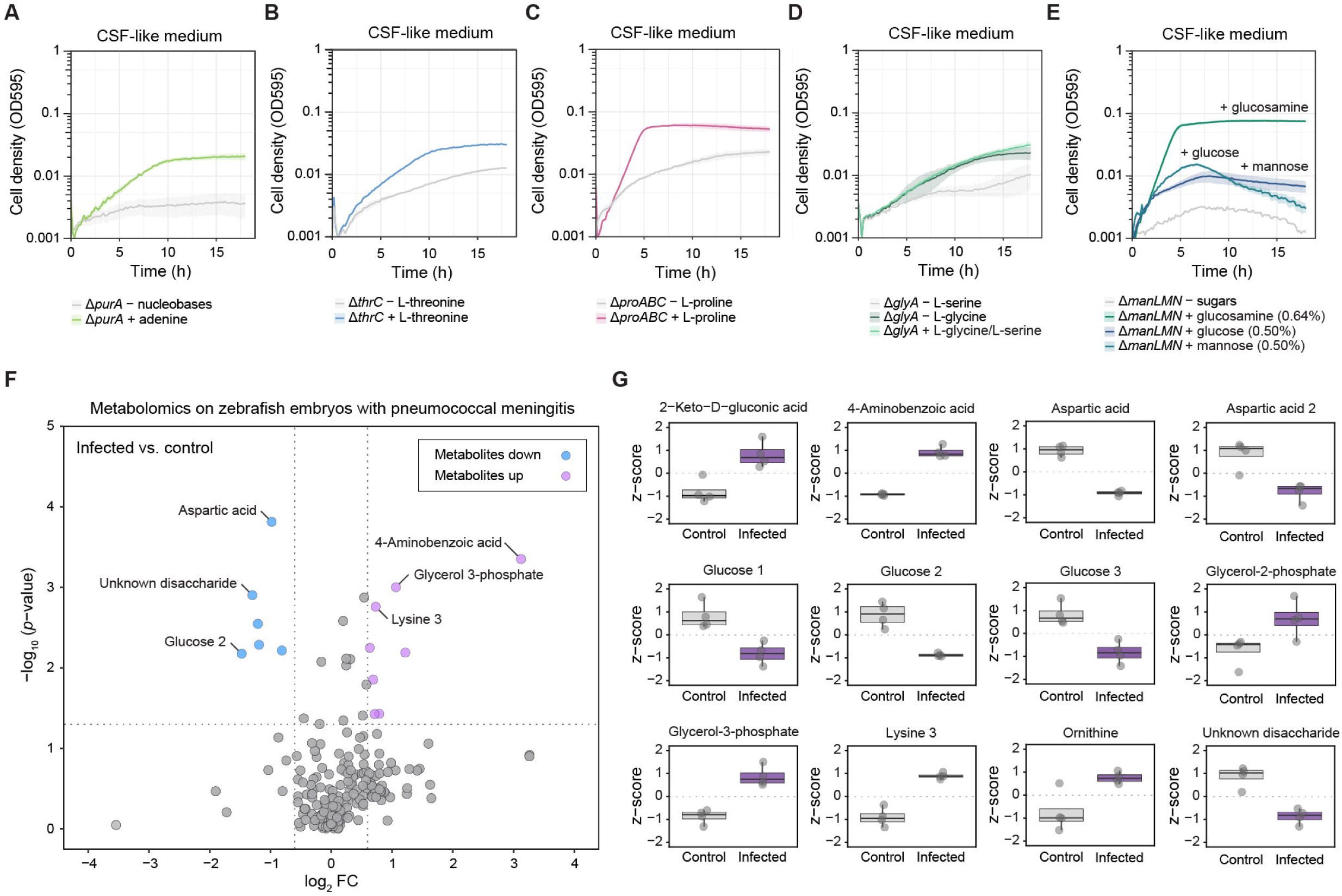
*In vitro* validation of metabolic mutants in cerebrospinal fluid (CSF)-like medium and *in vivo* whole-animal metabolomic profiling during pneumococcal meningitis. (A-E) Growth of *Streptococcus pneumoniae* D39V deletion mutant strains, previously implicated in metabolic pathways based on literature, tested in CSF-like medium under nutrient-deprivation or supplementation conditions. (A) D39V Δ*purA* mutant grown in medium without nucleobases or with supplementation of adenine. (B) D39V ΔthrC mutant grown without L-threonine or with L-threonine supplementation. (C) D39V ΔproABC mutant grown without L-proline or with L-proline supplementation. (D) D39V ΔglyA mutant grown without L-serine or L-glycine compared with L-serine and L-glycine supplementation. (E) manLMN mutant grown in medium without sugars or supplemented with glucosamine (0.64%), glucose (0.5%), or mannose (0.5%). Data represent mean ± SEM of three technical replicates. (F) Volcano plot of whole-animal metabolomics comparing D39V wild-type infected and uninfected control zebrafish embryos. Embryos were infected at 2 days post-fertilisation (2 dpf), and samples were collected at 24 hours post-infection (hpi). A total of 200 embryos per condition (four biological replicates of 50 embryos each) were analysed. Significance thresholds: *p*-value < 0.05 and |log₂FC| > 1. (G) Z-score analysis of selected metabolites differing between infected and control groups.

To link the bacterial fitness phenotypes observed *in vivo* to host metabolic conditions, we performed untargeted whole-animal metabolomic profiling of zebrafish embryos infected with *S. pneumoniae* D39V under approximately the same experimental conditions as the CRISPRi-Seq screen (i.e. without doxycycline induction). The aim was to determine whether infection triggers host-side metabolic changes that could explain the bacterial dependencies identified by CRISPRi-Seq. As shown in Figure 4F and Table S4, the global metabolomic comparison revealed a focused remodelling of host-associated metabolites during infection, particularly within carbohydrate, amino acid, and lipid pathways. Of the main changes in host metabolites, glucose was strongly depleted, consistent with enhanced pneumococcal carbohydrate use and the well-established clinical observation of hypoglycorrhachia in bacterial meningitis^3^ (Figure 4G).

In contrast, 4-aminobenzoic acid (PABA), a precursor for bacterial folate biosynthesis feeding one-carbon metabolism, was elevated, in line with prior work linking pneumococcal PABA/folate biosynthesis to growth and virulence under nutrient-restricted conditions^37^. Elevation of glycerol-3-phosphate and glycerol-2-phosphate suggests intensified phospholipid turnover and membrane remodelling during infection, which is consistent with host inflammatory responses and pneumococcal adaptation to lipid-derived carbon sources during invasive disease^38^. Both lysine and ornithine accumulated in CSF, indicating altered nitrogen flux and signalling through polyamine biosynthesis pathways that regulate both bacterial virulence and immune cell activation^39,40^. Additional metabolite shifts supported these core findings. Aspartic acid was reduced, consistent with its consumption as a critical substrate for both bacterial and immune cell nucleotide biosynthesis during infection^41^. The accumulation of organic acid metabolites, including 2-keto-D-gluconic acid and other glucose oxidation intermediates, reflects enhanced carbohydrate turnover during meningitis, including pyruvate oxidase-dependent H₂O₂ generation that drives both bacterial virulence and host inflammatory responses^42,43^. The overall pattern indicated limited systemic alterations, supporting that *S. pneumoniae* relies primarily on endogenous biosynthesis rather than major host-driven nutrient changes^36,44^. Baseline CSF metabolite concentrations are intrinsically low compared with plasma; thus, even modest bacterial consumption would not measurably deplete many amino acids or nucleobases such as adenine, proline, threonine, glycine, or serine^45^.

Together, the observed metabolite changes present in the host during infection mirror the pathways identified as essential in our CRISPRi-Seq screen, highlighting that *S. pneumoniae* must rely on de novo biosynthesis and efficient nutrient acquisition to persist in the nutrient-restricted CSF during meningitis.

### Defences against oxidative stress are essential for pneumococcal survival during meningitis

Our *in vivo* CRISPRi-Seq analysis identified nox and dpr as significantly more essential during pneumococcal meningitis, indicating that *S. pneumoniae* must withstand pronounced oxidative stress within the CSF. Further inspection of the *in vivo* dataset revealed that *trxB* (thioredoxin reductase) and *spxB* (pyruvate oxidase) also exhibited reduced fitness during infection, suggesting that pneumococci rely on multiple coordinated mechanisms to maintain redox balance *in vivo*.

During infection, *S. pneumoniae* encounters reactive oxygen species (ROS) from both its own metabolism and host immune responses (Figure 5A). Endogenously, hydrogen peroxide (H₂O₂) is produced by SpxB and lactate oxidase, whereas host-derived ROS are generated by the NADPH oxidase complex (NOX2/NCF1) during the neutrophil respiratory burst^46,47^. To counter oxidative stress, pneumococci employ NADH oxidase (Nox), which reduces O₂ while regenerating NAD⁺, and thioredoxin reductase (TrxB), which maintains the thioredoxin-TpxD system for detoxifying H₂O₂^11^. SpxB links central metabolism to peroxide production by converting pyruvate into acetyl phosphate, CO₂, and H₂O₂^48^.

**Figure 5.**
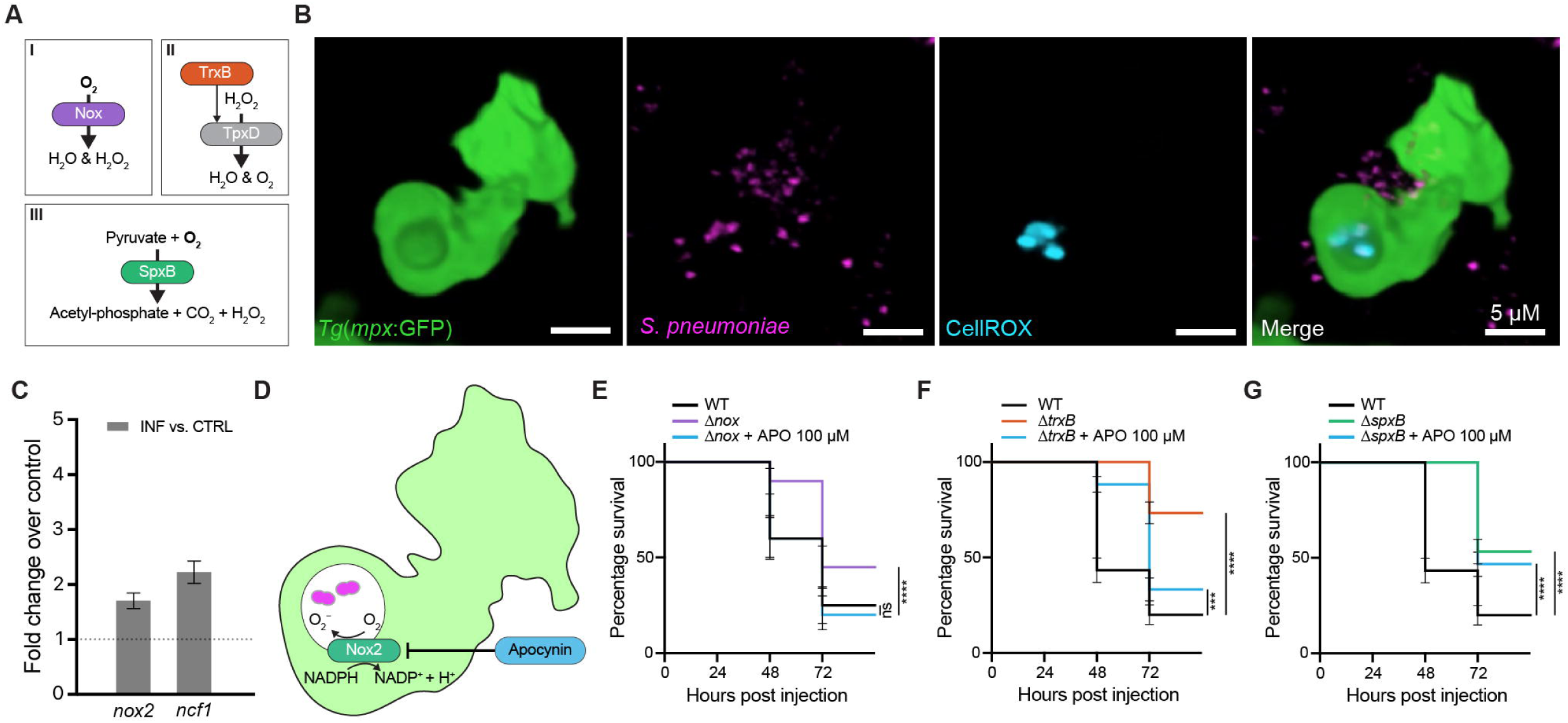
Pneumococcal and host oxidative-stress defence mechanisms shape infection outcome during meningitis. (A) Schematic overview illustrating how *S. pneumoniae* balances endogenous and host-derived oxidative stress through coordinated redox metabolism. The diagram highlights three *in vivo* essential genes identified in the CRISPRi-seq screen: *nox* (NADH oxidase) converts O₂ to H₂O and H₂O₂; *trxB* (thioredoxin reductase) maintains the thioredoxin system driving H₂O₂ detoxification via TpxD; and *spxB* (pyruvate oxidase) generates acetyl phosphate, CO₂, and H₂O₂ from pyruvate, linking central metabolism to peroxide production. Together, these enzymes maintain redox balance and support pneumococcal survival during oxidative stress *in vivo*. (B) Visualisation of ROS formation inside neutrophil phagolysosomes. *In vivo* live imaging of 2 days post-fertilisation (2 dpf) zebrafish embryos of the *Tg(mpx:*GFP) line infected with mScarlet-labelled *S. pneumoniae* D39V (VL1780). Approximately 1,000 CFU were injected into the hindbrain ventricle. CellROX Deep Red staining was used to detect ROS within neutrophil phagolysosomes during time-lapse confocal microscopy. The image shown was acquired at 2 hours post-infection (hpi) and represents a 3D-reconstructed view of the maximum projection of a z-stack. (C) Host oxidative-stress response during pneumococcal meningitis. Quantitative PCR analysis of (2 dpf) zebrafish embryos infected with *S. pneumoniae* D39V wild type (WT) or the D39 Δ*nox* knockout mutant. Embryos were collected at 6 hours post-infection (hpi) to assess host expression of nox2 and ncf1. Each condition included 20 embryos per replicate, with three biological replicates (n = 60 per group). (D) Schematic representation of NOX2 inhibition by apocynin, which prevents assembly of the NADPH oxidase complex and prevents ROS generation. (E-G) Survival analysis of infected zebrafish embryos treated with 100 µM apocynin. Embryos were infected at 2 dpf with approximately 300-400 CFU of the indicated pneumococcal D39V mutants and monitored for survival for up to 72 hpi. (E) D39V Δ*nox* deletion mutant is attenuated; apocynin treatment restores virulence phenotype. (F) D39V Δ*spxB* deletion mutant is attenuated; apocynin treatment does not restore virulence. (G) D39V Δ*trxB* deletion mutant is attenuated; apocynin treatment partially restores virulence. Each experiment represents three biological replicates with 20 embryos per group (n = 60 per condition). Data represent mean ± SEM. Statistical analysis was performed using the log-rank (Mantel-Cox) test; *p*-value < 0.05 was considered statistically significant: ns, not significant; *, *p*-value < 0.05; ***, *p*-value < 0.001; ****, *p*-value < 0.0001.

To confirm that reactive oxygen species (ROS) are generated during infection, zebrafish embryos of the *Tg(mpx:*GFP) line expressing green fluorescently labelled neutrophils were infected with mScarlet-labelled *S. pneumoniae* D39V (VL1780). CellROX, a fluorescent probe that emits upon oxidation and enables visualisation of ROS in live tissues, was directly injected into the hindbrain ventricle immediately after infection, revealing ROS accumulation within neutrophil phagolysosomes and indicating active oxidative responses during bacterial uptake (Figure 5B). Quantitative PCR of infected embryos at 24 hpi showed significant upregulation of *nox2* and *ncf1* compared with uninfected controls (Figure 5C), confirming activation of the host NADPH oxidase complex during pneumococcal meningitis.

To assess the bacterial contribution to oxidative stress resistance, we compared the virulence of Δ*nox*, Δ*trxB*, and Δ*spxB* mutants in the zebrafish embryo meningitis model. Although *dpr* was identified as more essential *in vivo*, it was excluded due to impaired *in vitro* growth. All three mutants were significantly attenuated compared with wild type (Figure 5E-G). Treatment with the NADPH oxidase inhibitor apocynin restored the virulence of Δ*nox* and partially rescued Δ*trxB*, whereas Δ*spxB* remained unaffected. These findings suggest that *nox* and *trxB* primarily protect against host-derived oxidative stress, while *spxB* contributes to endogenous redox metabolism and energy generation. Together, these results indicate that pneumococcal survival in the CSF depends on balancing protection against host-derived oxidative stress with regulation of its own metabolic ROS.

### Potential new therapeutic targets for pneumococcal infection

Antimicrobial resistance (AMR) in *S. pneumoniae* remains a major global concern, as resistance to β-lactams, macrolides and fluoroquinolones continues to complicate the treatment of invasive pneumococcal disease^49–51^. To explore alternative therapeutic strategies, we re-analysed our *in vivo* CRISPRi-Seq dataset and mapped genes with increased essentiality during pneumococcal meningitis to known and putative antibiotic-target pathways (Table S5).

Genes with increased essentiality were distributed across several key antibiotic-target pathways, including cell-wall synthesis, DNA and RNA synthesis, and ribosomal protein synthesis, as well as a distinct group of aminoacyl-tRNA synthetases (aaRSs) (Figure 6A). Within the cell-wall synthesis category, *pbp1a*, *pbp2b*, *pbp2x* and *pbp3* encode penicillin-binding proteins that represent classical β-lactam targets^52^. The ligases *ddl* and *murF*, involved in D-Ala-D-Ala dipeptide formation and peptide cross-linking of peptidoglycan, correspond to glycopeptide (vancomycin) targets^53^, while *uppS* and *ftsZ* are linked to cell-envelope assembly and division^54,55^. Genes related to nucleic acid synthesis included *gyrA*, *gyrB* and *parE* (fluoroquinolone targets)^56^, *rpoB* (the rifamycin target encoding the RNA polymerase β-subunit)^57^, and *polC* (encoding DNA polymerase III α-subunit)^58^. Ribosomal targets included the 50S subunit genes *rrlA-D* (23S rRNA) and *fusA* (elongation factor G), as well as the 30S subunit genes *rrsA-D* (16S rRNA) and *rpsL* (ribosomal protein S12), which are associated with macrolide, linezolid, tetracycline, and aminoglycoside activity^59^.

**Figure 6.**
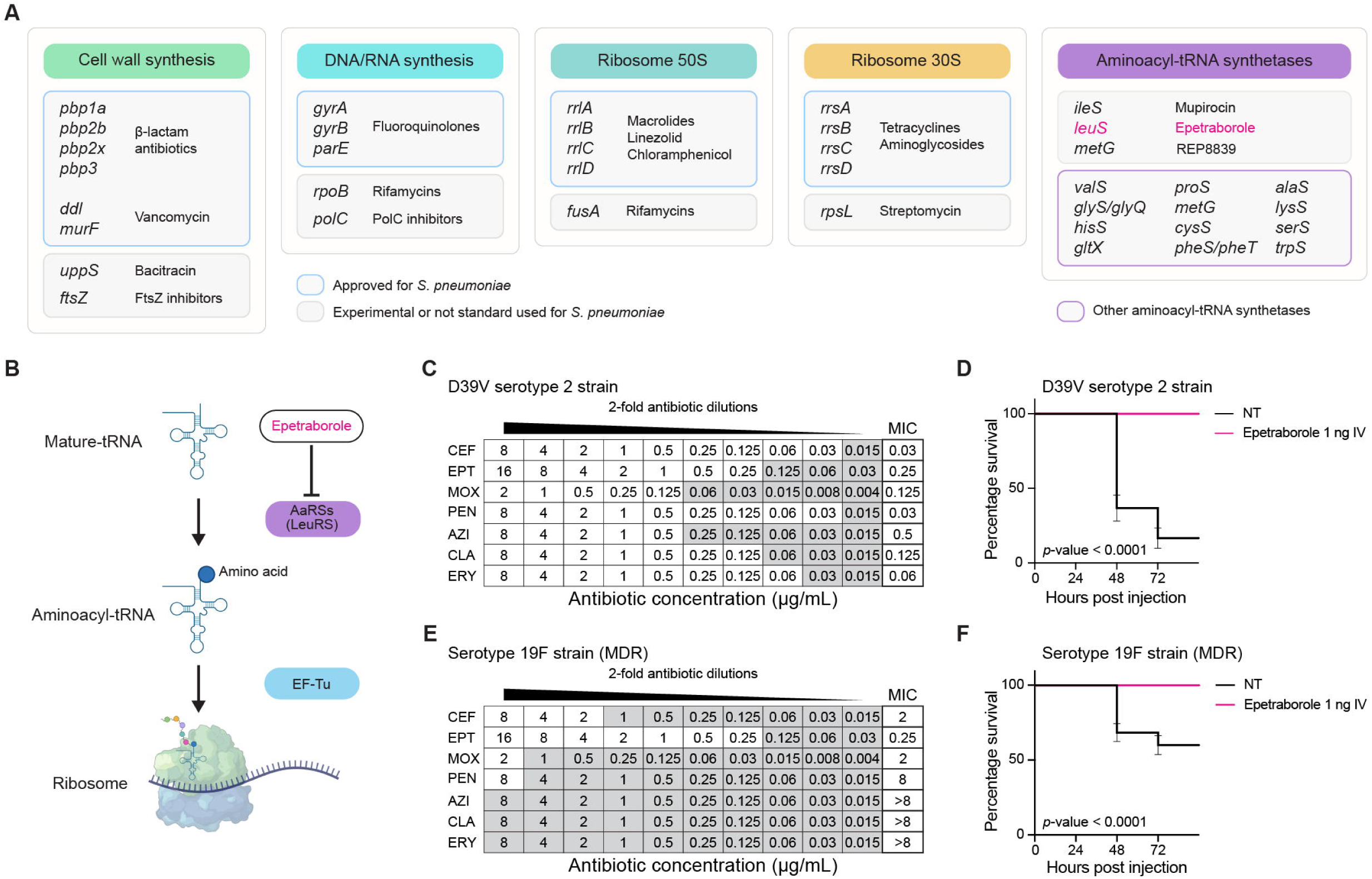
Discovery of aminoacyl-tRNA synthetases as novel *in vivo* drug targets in *Streptococcus pneumoniae* identified by CRISPRi-seq. (A) Re-analysis of the *in vivo* CRISPRi-seq dataset identified genes associated with established antibiotic targets, including pathways for cell-wall synthesis, DNA/RNA synthesis, and ribosomal 30S and 50S subunits. A distinct cluster of aminoacyl-tRNA synthetases (aaRS) also showed reduced fitness during infection. Genes were categorised as (blue) targets of approved antibiotics for *S. pneumoniae*, (grey) experimental or not routinely used agents, and (magenta) other aaRS-encoding genes. (B) Schematic representation of the boron-based antibiotic epetraborole, which inhibits leucyl-tRNA synthetase (LeuRS) by binding to its editing domain and preventing aminoacylation of tRNA. AaRSs, aminoacyl-tRNA synthetases; leucyl-tRNA synthetase (LeuRS); EF-Tu, elongation factor Tu. (C) Minimum inhibitory concentration (MIC) determination for *S. pneumoniae* D39V against ceftriaxone (CEF), epetraborole (EPT), moxifloxacin (MOX), penicillin (PEN), azithromycin (AZI), clarithromycin (CLA), and erythromycin (ERY), performed according to EUCAST microdilution protocols. (D) *In vivo* efficacy of EPT in the zebrafish embryo meningitis model. Embryos were infected at 2 days post-fertilisation (dpf) with approximately 300-400 CFU of D39V and treated with 1 nL of 1 mg/mL EPT at 1 hour post-infection (hpi). Each experiment represents three biological replicates with 20 embryos per group (n = 60 per condition). Data represent mean ± SEM. Statistical analysis was performed using the log-rank (Mantel-Cox) test; *p*-value < 0.05 was considered statistically significant. (E) MIC determination for a clinical multidrug-resistant (MDR) *S. pneumoniae* isolate (serotype 19F) resistant to macrolides, fluoroquinolones and β-lactams, assessed for the same antibiotics as in (C). (F) Survival analysis of zebrafish embryos infected with the 19F strain and treated with EPT as in (D). Each experiment represents three biological replicates with 20 embryos per group (n = 60 per condition). Data represent mean ± SEM. Statistical analysis was performed using the log-rank (Mantel-Cox) test; *p*-value < 0.05 was considered statistically significant.

Beyond these classes, a cluster of aminoacyl-tRNA synthetase (aaRS) genes also showed reduced fitness during infection. In total, 17 of 21 aaRS genes (81%) and 17 of 244 tRNA genes (6.9%) were significantly more essential *in vivo* (Table S1B). These enzymes catalyse the attachment of amino acids to their cognate tRNAs, a fundamental step in translation and protein synthesis^60^. Together, these findings define the *in vivo* antibiotic-target landscape of *S. pneumoniae* during meningitis, encompassing classical pathways of cell-wall, nucleic-acid, and protein synthesis, as well as aaRSs that represent a previously underexplored vulnerability.

Within this cluster, leucyl-tRNA synthetase (LeuRS) emerged as one of the most depleted genes *in vivo*. LeuRS catalyses the attachment of leucine to tRNA^Leu and is inhibited by the benzoxaborole epetraborole (EPT), which binds the enzyme’s editing domain and prevents aminoacylation^61^. EPT has demonstrated potent activity against Gram-negative bacteria and *Mycobacterium abscessus*^62–65^, but its potential against *S. pneumoniae* has not yet been examined.

To test this, we determined the minimum inhibitory concentration (MIC) of EPT against the *S. pneumoniae* D39V strain together with standard antibiotics (ceftriaxone, penicillin, moxifloxacin, azithromycin, clarithromycin, and erythromycin) using EUCAST microdilution protocols. EPT inhibited pneumococcal growth at 0.25 µg/mL (Figure 6C). Since no clinical breakpoints are yet available, this value should be interpreted as an indication of relative potency. In a zebrafish meningitis model, embryos infected with approximately 300-400 CFU of D39V and treated with 1 nL of 1 mg/mL EPT at 1 hour post-infection showed significantly improved survival compared with vehicle controls (Figure 6D). A similar effect was observed for a multidrug-resistant (MDR) clinical isolate (serotype 19F) resistant to β-lactams, macrolides, and fluoroquinolones, which showed the same apparent MIC (0.25 µg/mL) and improved survival *in vivo* (Figure 6E–F).

Together, these findings identify LeuRS as a critical *in vivo* target during pneumococcal meningitis and demonstrate that pharmacological inhibition of tRNA charging by EPT effectively restricts pneumococcal growth and improves host survival.

## Discussion

Our *in vivo* CRISPRi-Seq approach provides an unprecedented view of genes required for pneumococcal survival during meningitis. By systematically repressing nearly all pneumococcal genes *in vivo*, we identified 244 loci whose knockdown reduced bacterial fitness in the CSF. As expected, classical virulence determinants, including the polysaccharide capsule (*cps2A-cps2N* operon), were critical for survival during meningeal infection, validating the physiological relevance of our model^14^. Beyond these established factors, we also uncovered previously unrecognised genes involved in metabolism, cell-envelope biogenesis, and stress adaptation that become specifically required in the context of meningitis. These findings reveal how *S. pneumoniae* must reprogram its metabolic and defensive networks to persist within the infected CNS. Together, they define a meningitis-specific essentialome that integrates both known virulence factors and newly identified loci, and they expose promising therapeutic vulnerabilities such as aminoacyl-tRNA synthetases, including LeuRS, which can be targeted pharmacologically by the benzoxaborole epetraborole.

A key finding from the *in vivo* CRISPRi-Seq screen is that pneumococcal survival during meningitis depends on major metabolic adaptations. Deletion of *purA*, *proABC*, *thrC*, and *glyA*, genes involved in purine and amino acid biosynthesis, strongly reduced bacterial fitness *in vivo*. Growth of the corresponding mutants was rescued by adenine, proline, threonine, or serine, indicating that the cerebrospinal fluid (CSF) is nutrient-limited. CSF amino acid levels are intrinsically low compared with plasma due to restricted transport across the blood-brain barrier, creating a nutrient-poor environment that constrains bacterial growth^66^. Transcriptomic profiling of *S. pneumoniae* in CSF further showed strong induction of nucleotide and amino-acid biosynthetic pathways, consistent with adaptation to nutrient limitation^67^. These conditions make *de novo* purine and amino-acid synthesis pathways conditionally essential. Consistently, PurA was previously shown to be required for pneumococcal growth in human CSF and virulence in meningitis and pneumonia in murine models^25,32^, and disruption of *proABC* or one-carbon metabolism genes impairs CSF growth and systemic infection^68^ (Ramos-Sevillano et al., 2024). Together, these findings highlight nutrient limitation as a key selective pressure shaping pneumococcal metabolism during CNS infection.

Repression of the *manLMN* phosphotransferase system reduced pneumococcal fitness *in vivo*, and the strong rescue by glucosamine indicates that access to alternative hexoses supports persistence in the meningeal niche. ManLMN is a high-flux PTS permease for glucose, mannose, and glucosamine and also coordinates central carbon metabolism, so its down-regulation predictably limits growth when hexose availability is constrained^34,69^. Clinically, bacterial meningitis is characterised by low CSF glucose, reflecting a carbon-poor environment and heightened metabolic stress; elevated CSF lactate is also typical^3,70,71^. In this carbon-limited environment, *S. pneumoniae* maintains growth by exploiting alternative sugars through versatile uptake systems such as ManLMN^34,69^. Recent multi-omics data further suggest atypical sugar utilisation programs in pneumococci and link carbon metabolism to membrane homeostasis, suggesting that flexible carbohydrate uptake is a key selective advantage during infection^33^. These pathways could form the basis for selective therapeutic strategies by exploiting bacterial-specific vulnerabilities in metabolic regulation in the future.

A second major requirement for pneumococcal survival during meningitis is protection against oxidative stress. We identified several redox-associated genes (*nox*, *trxB*, *spxB*, and *dpr*), whose repression markedly reduced bacterial fitness and attenuated virulence *in vivo*, indicating that redox homeostasis is critical in the infected brain. Oxidative pressure arises from both bacterial and host sources: pneumococcal SpxB-dependent hydrogen peroxide production contributes to inflammation and virulence, while neutrophils and microglia generate reactive oxygen species through the NADPH oxidase complex as part of the host defence^11,42,48,72^. To maintain redox balance under these oxidative conditions, pneumococci rely on enzymes such as Nox, which detoxify molecular oxygen and regenerate NAD⁺ for central metabolism. Loss of Nox increases susceptibility to oxidative stress and markedly attenuates virulence in a mouse sepsis model^29^. Similarly, TrxB maintains the cellular thiol-redox balance and is required for oxidative-stress resistance and virulence^73,74^. In our model, deletion of nox or trxB attenuates virulence and increases ROS sensitivity. Inhibition of host NADPH oxidase with apocynin fully rescued the Δ*nox* but only partially restored the Δ*trxB* phenotype, indicating that Nox primarily counteracts host-derived oxidative stress, whereas TrxB also sustains intracellular redox homeostasis. Together, these findings highlight that oxidative stress defences are essential for pneumococcal pathogenesis, including during CNS infection. Because these bacterial systems overlap with host redox biology, future strategies could target pathogen-specific oxidative mechanisms while preserving physiological ROS signalling in the brain.

Beyond metabolic and stress adaptation, we also uncovered a distinct cluster of genes corresponding to known and putative antibiotic targets. A large subset encoded aminoacyl-tRNA synthetases (aaRS), several of which showed reduced fitness *in vivo*, suggesting that protein synthesis becomes a critical vulnerability during meningitis. Within this group, *leuS* was selected for follow-up because it can be specifically inhibited by the benzoxaborole epetraborole, which binds the enzyme’s editing domain and blocks aminoacylation^61^. *In vitro*, epetraborole inhibited pneumococcal growth at sub-microgram concentrations, and treatment in the zebrafish embryo meningitis model improved host survival and reduced bacterial burden. A MDR clinical isolate (serotype 19F) displayed comparable sensitivity, indicating that benzoxaboroles act independently of classical resistance mechanisms affecting β-lactams, macrolides, or fluoroquinolones.

While resistance to epetraborole has been reported in Gram-negative bacteria such as *Escherichia coli* and *Pseudomonas aeruginosa*, pneumococci differ in several key aspects that may favour sustained susceptibility^75^. The lack of a complex outer-membrane barrier, limited efflux pump repertoire compared to Gram-negative bacteria, and the essential, non-redundant nature of most aminoacyl-tRNA synthetases may limit opportunities for resistance development^76–78^. Nevertheless, recent structural work in *Staphylococcus aureus* demonstrated that mutations in the conserved HIGH motif of class I aaRS enzymes can confer extreme mupirocin resistance without loss of catalytic efficiency, underscoring that active site adaptation remains a potential route for Gram-positive aaRS resistance^79^. iven that multiple aaRS genes became essential during pneumococcal infection, this enzyme family represents an attractive and druggable target class for antimicrobial development. Notably, simultaneous inhibition of distinct aaRS enzymes was shown to markedly reduce the emergence of resistance^80^, underscoring their potential for combination or multitarget antimicrobial strategies^81^. Future work should combine structure-guided design of pneumococcal aaRS inhibitors with systematic evaluation of resistance trajectories under selective pressure and optimisation of next-generation benzoxaboroles for improved stability and pharmacokinetic properties. Together, these efforts could establish aaRS inhibition as a promising therapeutic strategy against multidrug-resistant pneumococci and other Gram-positive pathogens.

Despite the insights gained, several limitations should be considered. The zebrafish embryo model provides a transparent and tractable system for *in vivo* functional genomics, yet it differs from mammalian meningitis in temperature, immune maturation, and cerebrovascular physiology. These factors may influence bacterial fitness requirements and host-pathogen interactions. CRISPRi knockdown reduces, but does not completely abolish, gene expression, and polar effects on downstream genes cannot be excluded. Moreover, the screen was conducted at a single infection stage and inoculum, meaning that genes important for later or persistent phases of infection may not have been detected. Finally, the experiments were performed in one laboratory strain (D39V), and essentiality patterns are likely to vary among clinical isolates and serotypes. Addressing these aspects in mammalian meningitis models and across a broader strain panel will help define a core *in vivo* essentialome and clarify strain-specific adaptations.

In conclusion, this study provides the first comprehensive *in vivo* fitness map of *S. pneumoniae* during meningitis. By systematically repressing nearly all bacterial genes in the infected host, we uncovered a meningitis-specific essentialome that integrates classical virulence factors with newly identified metabolic, redox, and translational dependencies. These findings illustrate how *S. pneumoniae* adapts its physiology to survive within the nutrient-poor and oxidatively hostile environment of the cerebrospinal fluid. Beyond advancing our understanding of pneumococcal pathogenesis, this work highlights infection-specific vulnerabilities that can be exploited for therapeutic development, from metabolic inhibitors to aminoacyl-tRNA synthetase-targeting compounds such as benzoxaboroles. Together, these insights position *in vivo* CRISPRi-Seq as a powerful platform for dissecting bacterial gene function under physiological conditions and for guiding the rational discovery of new interventions against pneumococcal meningitis.

## Acknowledgements

We thank Louise Martin, Afonso Bravo, Johann Mignolet and Clement Gallay for technical assistance, Pedro Aires for assistance with zebrafish husbandry and embryo care, and Prof. Mark van der Linden for providing the multidrug-resistant clinical isolate *S. pneumoniae* serotype 19F strain. Selected illustrations were created with Biorender.com.

## Methods

### Bacterial strains and growth conditions

All pneumococcal strains used in this study are derivatives of the clinical isolate *S. pneumoniae* D39 from the Veening lab (D39V)^27^, unless specified otherwise and are listed in Table S6. Oligonucleotides are listed in Table S6. *S. pneumoniae* was grown at 37°C on Columbia blood agar (Thermo Fischer Scientific; CM0331B) plates supplemented with 2% sheep blood (Thermo Fisher Scientific; SR0051E) or in C+Y medium^82^. For transformation, pneumococcal cells were grown in C+Y medium at 37°C to an OD595 of approximately 0.1. Subsequently, 100 ng/mL of synthetic CSP-1 (AnaSpec, Cat# AS-63779) was added, and the cells were incubated for 10 min at 37°C. DNA was added to the activated cells and incubated for 20 min at 30°C. Cells were then diluted 10 times in fresh C+Y medium and incubated for 1.5 h at 37°C. Transformants were selected by plating on Columbia agar blood plates supplemented with 2% (v/v) defibrinated sheep blood with the appropriate antibiotics. Antibiotic concentrations for selection used for *S. pneumoniae* were: chloramphenicol (cam) 4.5 mg/mL, erythromycin (ery) 0.25 mg/mL or kanamycin (kan) 250 mg/mL.

### Construction of knockout mutant strains

Knockout mutants were generated by replacing the gene of interest with a kanamycin resistance cassette (kanᴿ). The resistance marker was amplified from genomic DNA of the *S. pneumoniae* D39 *hexA*::eryᴿ strain (Veening lab collection) using primers OVL10757 and OVL10758. The upstream and downstream flanking regions of the target gene were amplified from *S. pneumoniae* D39V genomic DNA using primer pairs listed in Table S7. The three resulting fragments, i.e. upstream region, resistance cassette, and downstream region, were assembled using Golden Gate cloning with BsaI restriction sites. The assembled constructs were transformed into wild-type *S. pneumoniae* D39V as described above. Transformants were selected on Columbia blood agar supplemented with 2% (v/v) defibrinated sheep blood and 250 µg/mL kanamycin. Correct genomic integration was confirmed by colony PCR.

### Zebrafish husbandry and maintenance

Transparent adult *tra^b6/b6^;nac^w^*^2*/w*2^ (TraNac) mutant zebrafish and *Tg*(*mpx*:GFP)/TraNac zebrafish expressing green fluorescently labelled neutrophils were maintained at 26°C in aerated 5-L tanks with a 10/14 h dark/light cycle^83^. Zebrafish embryos were obtained by natural mating and were collected within the first hours post-fertilisation and were raised at 28°C in a temperature-controlled incubator in E3 medium (5.0 mM NaCl, 0.17 mM KCl, 0.33 mM CaCl2·2H2O, 0.33 mM MgCl2·6H2O) supplemented with 0.3 mg/L methylene blue. Infection experiments were performed in 2 days post-fertilisation (dpf) zebrafish larvae unless stated otherwise. At this stage, the sex differentiation is not yet complete. All procedures involving zebrafish embryos were conducted according to local animal welfare regulations. The breeding of zebrafish in authorised institutions is in full compliance with the Swiss law on animal research. All procedures related to zebrafish studies at UNIL comply with the Swiss regulations on animal experimentation (cantonVaud, licence number VD-H28; Animal Welfare Act SR 455 and Animal Welfare Ordinance SR 455.1).

### Zebrafish infection experiments

All *S. pneumoniae* D39V strains were grown in C+Y medium until mid-log phase (OD595 = 0.2-0.3), harvested by centrifugation (6,000 RPM for 10 min), washed with sterile PBS, harvested again by centrifugation, and finally resuspended in 0.25% (w/v) amaranth solution (Sigma-Aldrich; A1016) to aid visualisation of the injection process. Before injection, 2 dpf embryos were mechanically dechorionated if necessary and anaesthetised in 0.02% (w/v) Tricaine (Sigma-Aldrich, A5040). Embryos were randomly assigned to experimental groups. The bacterial suspension was then injected in a volume of approximately 1 nL into the hindbrain ventricle of zebrafish embryos at 2 dpf for survival and bacterial load experiments. The number of colony-forming units (CFU) per injection was determined by quantitative plating of the injection volume. After injection, the larvae were kept in 6-well plates containing egg water (60 mg/mL sea salts (Sigma-Aldrich; S9883) in demi water) at 28°C, and the mortality rate was determined by monitoring live and dead embryos at fixed time points between 24 and 72 hours post-injection (hpi). To assess the effect of reactive oxygen species inhibition in the host, i.e. zebrafish embryos, on survival, infected zebrafish larvae were treated orally with 100 mM apocynin (Sigma-Aldrich; 178385) in 1% DMSO or 1% DMSO alone (vehicle control) by adding the compound to egg water for the whole infection course with refreshment of the egg water and compound every 24 h. All experiments were performed in triplicate. Survival graphs were generated with GraphPad Prism 10.0 and analysed with the log-rank (Mantel-Cox) test. Results were considered significant at *p*-values < 0.05.

### Bacterial load

To determine the bacterial load at different time-points, infected zebrafish larvae were anaesthetised in 0.02% Tricaine (Sigma-Aldrich, Cat# A5040) in egg water, transferred to a 1.5 mL screwcap tube (1 larva per tube) filled with 1.0mmglass beads (Sigma-Aldrich, Z250473) to 25% capacity of the tubes’ volume, placed in a microvial rack, and violently shaken (3 times 10 s, 10 s interval) in a FastPrep®-24 5G bead beating grinder and lysis system (MP Biomedicals, 11600550) to disrupt the cells and tissues. Subsequently, serial dilution plating was performed on Columbia Blood Agar plates supplemented with 2% defibrinated sheep blood, 10 mg/L colistin sulfate and 5 mg/L oxolinic acid (COBA; Oxoid, SR0126), to inhibit growth of commensal bacteria in zebrafish. The plates were incubated O/N at 37°C and quantified the next day. All experiments were performed in duplicate. Scatterplots were generated with GraphPad Prism 10.0 and analysed with an unpaired t-test.

### CRISPRi-Seq screen in zebrafish embryos

Pooled *S. pneumoniae* CRISPRi-Seq sgRNA libraries (VL4049) were initially cultured in C+Y medium at 37°C until reaching an optical density of OD595 = 0.3. These precultures were then diluted 1:100 into fresh C+Y medium and incubated at 37°C until mid-log phase (OD595 = 0.2-0.3). Bacterial cells were harvested by centrifugation at 6,000 rpm for 10 minutes, and the supernatant was discarded. Pellets were resuspended in an appropriate volume of 0.25% (w/v) amaranth dye solution in PBS, adjusted both to aid visualisation during injection and to achieve the desired CFU concentration per injection. Approximately 1 nL of the bacterial suspension was microinjected into the hindbrain ventricle of 2-day post-fertilisation (dpf) zebrafish embryos. For each condition, 50 embryos per replicate were injected (n = 200 per group across 4 biological replicates). Following injection, embryos were incubated at 28°C for 24 hours in egg water containing either 500 ng/mL doxycycline (induced) or no doxycycline (non-induced, DMSO control), to activate or suppress dCas9 expression, respectively. At 24 hpi, infected zebrafish embryos were collected, and gDNA isolation was performed using the DNeasy Blood &Tissue Kit (QIAGEN, 69504) for DNA isolation.

### Library preparation and sequencing

Illumina sequencing libraries were prepared using a one-step PCR protocol with primers listed in Table S7. Genomic DNA (gDNA) isolated from *S. pneumoniae* was used as the template. During the 30-cycle PCR, the index 1 (N701-N712), index 2 (N501-N508), and Illumina adapter sequences were introduced simultaneously. The amplicon sequence structure is shown below:

AATGATACGGCGACCACCGAGATCTACACTAGATCGC(N501)TCGTCGGCAGCGTCAGATGTGTATAAGAGACAGCCATTCTACAGTTTATTCTTGACATTGCACTGTCCCCCTGGTATAATAACTATANNNNNNNNNNNNNNNNNNNNGTTTAAGAGCTATGCTGGAAACAGCATAGCAAGTTTAAATAAGGCTAGTCCGTTATCAACTTGAAAAAGTGGCACCGAGTCGGTGCTTTTTTTCTGTCTCTTATACACATCTCCGAGCCCACGAGACTAAGGCGA(N701)ATCTCGTATGCCGTCTTCTGCTTG

Here, N501 and N701 represent i5 and i7 barcode indices, respectively, and the highlighted 20 N region denotes the variable sgRNA base-pairing sequence. A full annotation of the amplicon design is available in the file “CRISPRi-seq amplicon.dna” on the Veening Lab website (https://www.veeninglab.com/crispri-seq). Each 50 µL PCR contained 4 µg of gDNA as template. The 304 bp amplicons were purified from 1 % agarose gels and quantified using a Qubit dsDNA assay (Thermo Fisher Scientific; Q32854). Purified amplicons were sequenced on an Illumina NovaSeq 6000 system using a custom run configuration: the first 54 cycles (common sequence) were dark cycles, followed by 20 cycles to read the variable sgRNA base-pairing region.

### sgRNA read mapping and quantification

Quantification of single guide RNA (sgRNA) abundances from the *S. pneumoniae* CRISPRi-seq screens during zebrafish infection was performed using 2FAST2Q (v2.5.5), a Python-based tool developed by the Veening Lab for rapid extraction of short, defined sequences such as sgRNAs or barcodes from large-scale sequencing datasets^84^. Raw FASTQ files were first demultiplexed by Illumina index, after which sgRNA sequences were extracted from read 1 starting at position 54, allowing for up to one mismatch against the reference library of 1,498 sgRNAs sequences used in the CRISPRi library. Reads with base quality scores < 24 at any sgRNA position were discarded. The following parameters were used:

--start 54 --length 20 --mismatch 1 --phred 24.

Resulting raw counts for all sgRNAs are provided in Table SX. The 2FAST2Q tool is freely available at https://github.com/veeninglab/2fast2q.

### Differential fitness analyses

To evaluate the fitness cost of each sgRNA during infection, count data were analysed using the DESeq2 package in R^85^. Differential abundance testing was performed against a |log₂| fold-change (log₂FC) threshold of 1, with an adjusted significance level of α = 0.05.

### Interaction analysis

Raw reads of the CRISPRi library grown in C+Y medium were downloaded from SRA with NCBI accession number PRJNA611488 ^25^. sgRNA counts were quantified as described above using 2FAST2Q. Differential enrichment analysis between the libraries derived from zebrafish and C+Y samples was performed using DESeq2 as described above. To account for any bias caused by a difference in the overall number of generations the bacteria grew in the presence of the CRISPRi inducer between the two conditions, we estimated generation numbers in the zebrafish model based on the bacterial load at 24 hpi. We incorporated this into the DESeq2 model design by scaling the induction generation numbers (21 for C+Y medium, 5 for zebrafish samples, and 0 for all uninduced samples regardless of medium) using the built-in R function scale(), and adding it to the model as an explanatory variable, along with the medium variable (C+Y medium or zebrafish-derived) and their interaction. This interaction term represents the differential sgRNA enrichment upon CRISPRi induction between the two growth conditions.

### Gene Ontology (GO) enrichment analysis

Gene Ontology (GO) enrichment analysis was conducted to identify overrepresented biological processes, molecular functions, and cellular components among genes classified as “more essential” or “more costly” (showing increased fitness upon CRISPRi knockdown) in *S. pneumoniae* D39V during meningitis. Input gene sets comprised either 435 unique essential genes or 134 unique costly genes identified from *in vivo* CRISPR interference-based fitness profiling. The reference background consisted of all 1,996 annotated protein-coding genes in the *S. pneumoniae* D39V genome (BioCyc version 29.0 and PneumoBrowse 2^87^. GO term annotations were compiled from BioCyc, PneumoBrowse, and eggNOG-mapper outputs to ensure maximal and accurate gene-GO mapping. For analysis, each gene was mapped to all associated GO terms in the categories Biological Process (BP), Molecular Function (MF), and Cellular Component (CC). For each GO term, a 2×2 contingency table was created to record the number of genes annotated with the term in both the input set and the genome-wide background. Enrichment for each GO term was assessed using a one-sided Fisher’s exact test (testing for overrepresentation in the input set compared to the background). For each GO category, raw *p*-values were adjusted for multiple hypothesis testing using the Benjamini-Hochberg false discovery rate (FDR) procedure. Enrichment was considered significant only if the FDR-adjusted *p*-value was below 0.05. All calculations were performed in Python (pandas, scipy.stats) and R (base and stats packages). Final enrichment tables were generated as CSV files, and figures were created in R using ggplot2. Only GO terms with robust annotation counts (≥3 genes per set) were reported. Care was taken to interpret results in the context of the hierarchical and correlated nature of the GO ontology, and biological conclusions were drawn only for terms surviving multiple-testing correction.

### Immunohistochemical staining

To visualise the ventricular lining and neuroepithelium of the zebrafish brain after pneumococcal infection, 2 dp) embryos were infected in the hindbrain ventricle with the dual fluorescent *S. pneumoniae* reporter strain VL2359 and collected at 24 hpi. Embryos were fixed overnight in 4% paraformaldehyde (PFA) in PBS, then stained with a zona occludens-1 (ZO-1) monoclonal antibody (Invitrogen, 33-9100) to label tight junctions of the ventricular epithelium. Fixed embryos were rehydrated in PBTx (1% Triton X-100 in PBS), permeabilised with 0.25% trypsin in PBS, and blocked for 10 min in blocking buffer to minimise non-specific binding. Embryos were incubated overnight at room temperature on a seesaw rocker with ZO-1 antibody (1:400 v/v) diluted in antibody buffer (1% normal goat serum + 1% BSA in 1% PBTx). After washing and re-blocking, embryos were incubated overnight at 4 °C with Alexa Fluor 647 goat anti-mouse IgG (Life Technologies, A21235; 1:400 v/v).

### Confocal imaging of fixed samples

After immunohistochemical staining, embryos were mounted in 0.5% low-melting-point agarose (Thermo Fisher Scientific, R0801) in PBS and imaged in open, uncoated 8-well µ-slides (Ibidi). Confocal imaging was performed using a Nikon AXR resonant-scanning confocal microscope equipped with a Nikon Ti2 body, Okolab temperature-controlled chamber, and Hamamatsu ORCA-Flash 4.0 V3 camera. Image acquisition was performed with NIS-Elements AR (v5.42.06, Nikon), and images were processed using ImageJ (Fiji) for brightness and contrast adjustment, and visualisation of bacterial localisation relative to the ventricular epithelium.

### In vivo confocal imaging of reactive oxygen species (ROS)

For live imaging of reactive oxygen species (ROS) and host-pathogen interactions, *Tg(mpx:*GFP*)/TraNac* zebrafish embryos at 2 dpf were infected in the hindbrain ventricle with the red fluorescently labelled *S. pneumoniae* strain D39V *hlp*A::*hlpA_hlpA*-mScarlet-I (VL1780)^88^. Immediately after bacterial injection, 1 nL of CellROX™ Deep Red reagent (Thermo Fisher Scientific, C10422) was co-injected into the same site to enable *in vivo* detection of ROS. Before confocal imaging, infected embryos were screened for fluorescent neutrophils using a Nikon SMZ stereomicroscope equipped with a Nikon DS-Qi2 camera; only embryos showing robust *mpx*:GFP expression were selected. Embryos were embedded in 0.5% low-melting-point agarose containing 0.02% (w/v) Tricaine (Sigma-Aldrich, A5040) dissolved in PBS and imaged in open, uncoated 8-well µ-slides (Ibidi). Time-lapse fluorescence images were acquired every 5 min for 3 h using the same Nikon AXR setup. Image processing was performed in NIS-Elements AR (v5.42.06, NIkon and ImageJ (Fiji) for contrast enhancement, channel merging, and localisation of ROS and bacterial-neutrophil interactions.

### Microtiter plate-based growth and rescue assays

For *S. pneumoniae* growth assays, cells were precultured in C+Y medium until reaching an OD595 = 0.1, then diluted 100-fold into 96-well microtiter plates containing fresh C+Y medium, supplemented with the appropriate concentrations of antibiotics, inducers, or metabolites as indicated. Plates were incubated at 37°C, and OD595 was recorded every 10 minutes for 15-18 h using a TECAN Infinite M200 Pro plate reader. Each condition was tested in technical triplicate. For metabolite rescue experiments, mutant strains were grown in CSF-like medium^35^ lacking the corresponding nutrient and supplemented, where indicated, with adenine (Δ*purA*), proline (Δ*proABC*), threonine (Δ*thrC*), serine or glycine (Δ*glyA*), or glucose, mannose, or glucosamine (ΔmanLMN). Growth was monitored continuously under identical conditions.

### RT-qPCR

For zebrafish gene expression analysis, total RNA was extracted from zebrafish embryos (n = 20 per biological replicate unless stated otherwise) at 24 hpi. Larvae were anaesthetised in 0.02% Tricaine (Sigma-Aldrich, A5040) in egg water, transferred to 2 mL microcentrifuge tubes, and the residual water was removed. Lysis buffer (Qiagen RNeasy Mini Kit; 74104) was added, and larvae were homogenised by repeatedly drawing the suspension through a 26-gauge needle attached to a 1 mL syringe until fully lysed. Total RNA was purified using the RNeasy Mini Kit (Qiagen; 74104) according to the manufacturer’s instructions, including on-column DNase digestion. Complementary DNA (cDNA) was synthesised from purified RNA using the High-Capacity cDNA Reverse Transcription Kit with RNAse Inhibitor (Thermo Fisher Scientific; 4374966) following the manufacturer’s protocol. Primers were designed across exon-exon junctions using NCBI Primer-BLAST and are listed in Table S6. Each quantitative PCR reaction contained *Power*SYBR^®^ Green PCR Master Mix (Thermo Fisher Scientific; 4367659), 0.5 µM of each primer, and 1 µL of cDNA template in a final volume of 10 µL. Reactions were performed in triplicate on a QuantStudi 5 Real-Time PCR System (Applied Biosystems) using standard cycling parameters. Expression levels were normalised to the housekeeping gene *bactin1* (β-actin-1), and relative expression changes were calculated using the ΔΔCt method^89^.

### Minimum inhibitory concentration (MIC) determination

Minimum inhibitory concentrations (MICs) were determined by broth microdilution according to EUCAST guidelines^90^. Assays were performed in cation-adjusted Mueller-Hinton broth supplemented with 5% lysed horse blood and 20 mg/L β-NAD (MH-F broth) in sterile 96-well microtiter plates. Bacterial suspensions were adjusted to a 0.5 McFarland standard and diluted to achieve a final inoculum of approximately 5 × 10⁵ CFU/mL per well. Plates were sealed and incubated at 37°C for 20 h. MICs were recorded as the lowest antibiotic concentration that completely inhibited visible growth, following the EUCAST Reading Guide for Broth Microdilution. The following antimicrobial agents were tested: cefotaxime, penicillin, moxifloxacin, azithromycin, clarithromycin, erythromycin, and epetraborole.

### In vivo epetraborole treatment

To evaluate the therapeutic potential of epetraborole *in vivo*, *TraNac* zebrafish embryos were infected at 2 dpf in the hindbrain ventricle with *S. pneumoniae* D39V wild-type or S. pneumoniae serotype 19F strain as described above. At 1 hpi, 1 nL of epetraborole (1 mg/mL in 0.25% amaranth in PBS) or vehicle (0.25% amaranth in PBS) was microinjected into the caudal vein for intravenous treatment. Embryos were maintained at 28°C in egg water throughout the experiment. Survival was monitored at fixed intervals up to 72 hpi. Each experiment included three biological replicates with 20 embryos per group (n = 60 per condition). Survival curves were generated in GraphPad Prism 10.0 and analysed using the log-rank (Mantel-Cox) test; *p*-value < 0.05 was considered statistically significant.

### Metabolite extraction and GC-MS analysis

For metabolomic profiling, zebrafish embryos were infected at 2 dpf with *S. pneumoniae* D39V wild-type or mock-injected as controls. At 24 hours post-infection (hpi), larvae were pooled in groups of 50 embryos per biological replicate (n = 4 per condition; total 200 embryos per group) and immediately processed for metabolite extraction. Soluble metabolites were extracted using a modified two-phase chloroform:methanol:water (Bligh and Dyer) protocol as previously described^91^. Briefly, pooled larvae were homogenised on ice, and metabolites were extracted in cold methanol and chloroform. L-norleucine (Sigma-Aldrich, N8513) and vanillic acid (Sigma-Aldrich, 94770) were added as internal standards prior to extraction. After centrifugation and phase separation, the aqueous fractions were collected and dried under vacuum. Dried samples were derivatised with 50 µL of 20 mg/mL methoxyamine hydrochloride in pyridine (Sigma-Aldrich, 226904) for 1.5 h at 33 °C, followed by 50 µL of MSTFA (Sigma-Aldrich, 69478) for 2 h at 45 °C. Immediately after derivatisation, 1 µL of each sample was injected by autosampler into an Agilent 8890/5977B GC-MSD system equipped with a VF-5MS column (30 m × 0.25 mm × 0.25 µm). The inlet operated in split mode (2:1) with a helium flow rate of 1 mL/min at 280 °C. The GC oven program was as follows: 125 °C (2 min hold), ramp 5 °C/min to 225 °C, then 15 °C/min to 310 °C (3.3 min hold). The mass spectrometer was operated in scan mode (50–550 Da) at 2.9 scans/s.

### Metabolomics data processing and statistical analysis

Raw GC-MS data were processed using MZmine2 (version 4.7.27)^92^ with the following modules: Scan-by-scan filtering, Mass detection, Chromatogram builder, Local minimum feature resolver, GC-EI spectral deconvolution, GC aligner, Feature list rows filter, Duplicate peak filter, Feature finder, and NIST MS Search (NIST 17 Mass Spectral Library). Metabolite annotations were accepted only for features with a cosine similarity > 0.7, and all hits were manually curated to confirm derivatised metabolite identities. Subsequent analysis was performed in R. Raw peak areas were normalised to the average response of the internal standards in each sample. Z-scores, fold changes, and (adjusted) *p*-values were calculated between control and infected groups using the Wilcoxon rank-sum test with Benjamini-Hochberg correction for multiple comparisons (Table S4).

### Statistical analysis

All statistical analyses were performed using GraphPad Prism 10.0, R (version 4.3.2) and RStudio (version 2025.05.1+513). Survival data were analysed using the log-rank (Mantel-Cox) test. Differences in bacterial load, metabolite abundance, or gene expression were assessed using unpaired t-tests or one-way ANOVA, as appropriate. For metabolomics, significance testing was performed using the Wilcoxon rank-sum test with Benjamini-Hochberg correction for multiple comparisons. Differential gene fitness from CRISPRi-seq screens was analysed in R using the DESeq2 package with an adjusted α = 0.05 and |log₂FC| ≥ 1 as the threshold for significance; effect-size shrinkage was applied using the apeglm method. Data were visualised with ggplot2 and GraphPad Prism 10.0. Unless otherwise specified, *p*-values < 0.05 were considered statistically significant.

### Data and code availability

Processed count matrices, analysis scripts, and supporting R code are available at https://github.com/veeninglab/CRISPRi-seq. Microscopy datasets, including raw confocal image files and representative videos of *in vivo* imaging, are available from the corresponding author upon reasonable request.

