## Supplementary figures and images for "*In vivo* genome-wide CRISPRi-Seq in *Streptococcus pneumoniae* reveals host-adaptive pathways essential for meningitis and identifies new therapeutic targets"

### Supplemental Figure 1

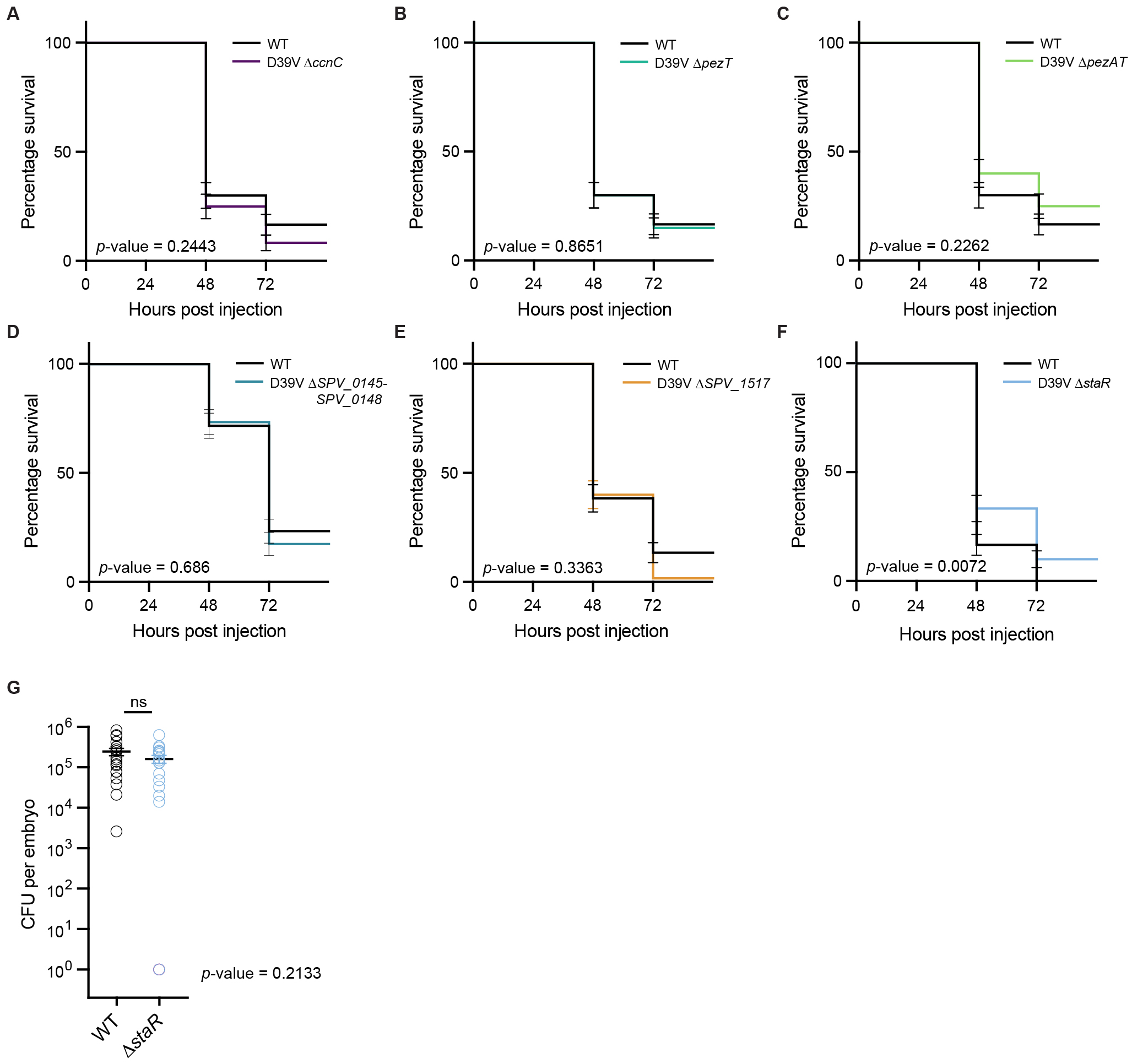
