## Supplementary Information for "*In vivo* genome-wide CRISPRi-Seq in *Streptococcus pneumoniae* reveals host-adaptive pathways essential for meningitis and identifies new therapeutic targets"

**Supplementary Figure 1. Survival analysis of zebrafish larvae infected with pneumococcal knockout mutants.**

(A-F) Survival of zebrafish larvae infected with *S. pneumoniae* D39V knockout mutants that did not show a significant difference in virulence compared to the wild type. Two days post-fertilisation (2 dpf) embryos were microinjected with 300-400 CFU of pneumococci into the hindbrain ventricle. Data represent the mean ± SEM of three independent biological replicates with 20 larvae per group (n = 60 per condition). Survival was analysed using the log-rank (Mantel–Cox) test; p < 0.05 was considered statistically significant. (A) *ccnC*, encoding a cell cycle–regulated protein. (B) *pezT*, encoding the toxin component of the PezAT toxin–antitoxin system. (C) *pezAT* operon, encompassing both toxin and antitoxin genes. (D) *staR*, encoding a competence-stimulating peptide regulator. (E) *SPV_1517*, encoding a putative transcriptional regulator. (F) *SPV_0145–0148*, encoding a predicted ABC transporter operon. (G) Bacterial load comparison between wild-type and *staR* mutant infections at 24 h post-infection. Data were analysed using the Mann–Whitney U test; p < 0.05 was considered statistically significant.

**Table S1. Differential fitness analysis during pneumococcal meningitis by CRISPRi-seq.** List of *S. pneumoniae* sgRNA targets from the *in vivo* CRISPRi-seq pneumococcal meningitis screen. Genes with significant depletion or enrichment are classified as essential, neutral or costly for bacterial survival during meningitis.

**Table S2. Gene Ontology (GO) enrichment analysis.** Summary of biological processes, molecular functions, and cellular components significantly enriched among genes identified as essential or costly in vivo. GO terms were determined using over-representation analysis with false-discovery-rate (FDR) correction.

**Table S3. Differential fitness analysis *in vivo* versus *in vitro***. Comparison of gene fitness profiles obtained from CRISPRi-seq during meningitis (*in vivo*) and in rich C+Y medium (*in vitro*).

**Table S4. Metabolomics analysis of infected versus control zebrafish embryos.** Untargeted GC-MS profiling on whole zebrafish embryos to compare metabolic signatures between *S. pneumoniae*-infected and uninfected controls.

**Table S5. Essential genes involved in antimicrobials.** List of genes encoding known or putative targets of antimicrobials identified as essential during meningitis.

**Table S6. Strains.** List of *S. pneumoniae* strains used in this study.

**Table S7. Primers.**  List of primers used in this study for cloning, qPCR, mutant construction, and validation.

**Table S8. Raw counts in vivo CRISPRi-seq.** Raw read counts for all sgRNAs recovered from the *in vivo* CRISPRi-seq screen during pneumococcal meningitis. Data represent sequencing reads per guide RNA across biological replicates before and after induction, forming the basis for the differential fitness analysis presented in Table S1.
